# Mate-finding dispersal reduces local mate limitation and sex bias in dispersal

**DOI:** 10.1101/770412

**Authors:** Abhishek Mishra, Sudipta Tung, V.R. Shree Sruti, Sahana V. Srivathsa, Sutirth Dey

**Affiliations:** Population Biology Laboratory, Biology Division, Indian Institute of Science Education and Research-Pune, Dr. Homi Bhabha Road, Pune, Maharashtra, India, 411 008

**Author notes:** Department of Organismic and Evolutionary Biology, Harvard University, Cambridge, Massachusetts, USA, 02138. McKnight Brain Institute, Life Sciences North Building, University of Arizona, Tucson, Arizona, USA, 85724. **Name and address of the corresponding author**: Sutirth Dey, Professor, Biology Division, Indian Institute of Science Education and Research, Pune, Dr. Homi Bhabha Road, Pune, Maharashtra, India 411 008.

**Keywords:** *Drosophila melanogaster*, dispersal propensity, temporal dispersal profile, dispersal plasticity, dispersal evolution, spatial sorting

## Abstract

1. Sex-biased dispersal (SBD) often skews the local sex ratio in a population. This can result in a shortage of mates for individuals of the less-dispersive sex. Such mate limitation can lead to Allee effects in populations that are small or undergoing range expansion, consequently affecting their survival, growth, stability and invasion speed.
2. Theory predicts that mate shortage can lead to either an increase or a decrease in the dispersal of the less-dispersive sex. However, neither of these predictions have been empirically validated.
3. To investigate how SBD-induced mate limitation affects dispersal of the less-dispersive sex, we used *Drosophila melanogaster* populations with varying dispersal propensities. To rule out any mate-independent density effects, we examined the behavioral plasticity of dispersal in presence of mates as well as same-sex individuals with differential dispersal capabilities.
4. In the presence of high-dispersive mates, the dispersal of both male and female individuals was significantly increased. However, the magnitude of this increase was much larger in males than in females, indicating that the former show greater mate-finding dispersal. Moreover, the dispersal of either sex did not change when dispersing alongside high- or low-dispersive individuals of the same sex. This suggested that the observed plasticity in dispersal was indeed due to mate-finding dispersal, and not mate-independent density effects.
5. Strong mate-finding dispersal can diminish the magnitude of sex bias in dispersal. This can modulate the evolutionary processes that shape range expansions and invasions, depending on the population size. In small populations, mate-finding dispersal can ameliorate Allee effects. However, in large populations, it can dilute the effects of spatial sorting.

## 1. Introduction

Dispersal, i.e. movement of organisms with potential consequences for gene flow (Ronce 2007), is a key life-history trait (Bonte & Dahirel 2017) that determines the temporal and spatial distribution of individuals. Dispersal has possible effects on population dynamics, biological invasions and community assembly (reviewed in Bowler & Benton 2005; Clobert *et al.* 2009; Lowe & McPeek 2014). Interestingly, in many sexual species, males and females can have very different dispersal properties. For example, males are known to exhibit higher dispersal in taxa such as lions (Packer & Pusey 1987), great white shark (Pardini *et al.* 2001) and spotted hyena (Höner *et al.* 2007). In contrast, many avian species typically exhibit a female bias in dispersal (reviewed in Clarke, Sæther & Roskaft 1997). Besides affecting the process of net dispersal itself, sex-biased dispersal (SBD) may have a pronounced bearing on phenomena such as adaptation to heterogeneous habitats (Kawecki 2003; Lopez *et al.* 2008), as well as invasions and range expansions (Miller & Inouye 2013). In sexually reproducing populations, the magnitude of SBD can also modulate the strength of Allee effects, which In turn may affect their survival, growth, stability and invasion speed (Taylor & Hastings 2005; Miller *et al.* 2011; Shaw & Kokko 2015). Given the academic as well as practical implications of these phenomena, SBD has been a major topic of investigation over the last several decades (reviewed in Pusey 1987; Clarke, Sæther & Roskaft 1997; Prugnolle & De Meeûs 2002; Lawson Handley & Perrin 2007).

A key question in the SBD literature has been the relationship between SBD and mating. Greenwood (1980) postulated that SBD occurs due to local competition for mates or other resources, and is further reinforced by the role of dispersal in inbreeding avoidance. As a result, he predicted that the direction of SBD is determined by the type of mating system (Greenwood 1980). The latter has often served as the conceptual framework for empirical studies on SBD across taxa (for example, see Ribble 1992; Langen 1996; Croft *et al.* 2003; Nagy *et al.* 2007; Pérez‐Espona *et al.* 2010). However, a recent analysis of data across several species suggested that evolution of SBD is better explained by sexual dimorphism and parental care (Trochet *et al.* 2016). Similarly, in a review of theoretical literature on the topic, Li and Kokko (2019) reported that the association between SBD and mating may not be limited to the type of mating system. Instead, the relative order of mating and dispersal in a given species might be a more important factor in this context (Li & Kokko 2019). However, while these studies discuss the role of mating as a cause of SBD, the consequences of SBD on mating have remained relatively unexplored.

Among other things, SBD can lead to a skew in the local sex ratio in the population. This, in turn, can affect the movement of the less dispersive sex in different ways, depending on the exact nature of the selection pressure faced by them (Shaw & Kokko 2014). For example, dispersal acts as a mechanism to avoid inbreeding in many species (Greenwood, Harvey & Perrins 1978; Pusey 1987; Lambin 1994). Under such circumstances, emigration of any one sex (say males) from the population could be sufficient for promoting mating between unrelated individuals, thus allowing the other sex (females in this example) to potentially reduce its investment in dispersal (Shaw & Kokko 2014; Fromhage, Jennions & Kokko 2016). This would result in even lower dispersal by the less-dispersive sex, thus, widening the magnitude of sex bias in dispersal. However, once the difference in dispersal between the sexes becomes high, it can lead to mate limitation and a reduction in local mating opportunities (Meier, Starrfelt & Kokko 2011; Miller *et al.* 2011; Miller & Inouye 2013). This could then lead to an increased pressure on the less-dispersive sex for a subsequent “mate-finding dispersal”, thus reducing the magnitude of difference in SBD (Meier, Starrfelt & Kokko 2011). Thus, there exists a ways for both, a decrease or an increase, in the dispersal of the less-dispersive sex following a SBD event.

A straightforward way for studying this phenomenon is to examine and compare dispersal under varied sex ratios. However, empirical studies on dispersal under artificially skewed sex ratios have either failed to observe SBD at all (Trochet *et al.* 2013), or found that the sex bias in movement was independent of the sex ratio (Baines, Ferzoco & McCauley 2017). Interestingly, these empirical studies investigated the effects of an existing skew in the sex ratios in the population. However, if one sex disperses more than the other does, then a skew in sex ratio can develop over the course of dispersal, even if the population had roughly equal proportions of the two sexes to begin with. To the best of our knowledge, the effect of this kind of a skew on the less-dispersive sex has never been investigated empirically.

Here, we investigate the interplay between dispersal and mate availability using laboratory populations of *Drosophila melanogaster* under controlled environmental conditions. Specifically, we asked the following questions: (1) is the movement of individuals affected by differential dispersal of mates? (2) Is the movement of individuals affected by differential dispersal of same-sex individuals? (3) Do males and females differ in their response in the above two scenarios? We focus on the fact that the sex ratio gets gradually skewed along the course of an SBD event, rather than abruptly, resulting in a time-dependent decrease in local mate availability. Therefore, we study the temporal dispersal profile of individuals in response to high-vs. low-dispersive mates. Moreover, we also study the dispersal of flies in response to high vs. low dispersive individuals of the same sex, to control for a potential confound of sex-independent density-dependent response. For these experiments involving mixed-sex and same-sex dispersal, we used individuals from fly populations with different dispersal properties. We found that neither male nor female dispersal was affected by the dispersal pattern of individuals of the same sex. However, the dispersal of mates significantly affected movement, more so in males than in females. We discuss the potential reasons for the observed behavioral plasticity, and its implications for SBD as well as dispersal evolution.

## 2. Materials and Methods

### 2.1 Fly populations

We used four different laboratory-maintained populations of *D. melanogaster* in this study (namely, DB_4_, DBS, VB_4_, and VBC_4_). All these populations are outbred and maintained at large population sizes (breeding size ~2400 individuals) at uniform environmental conditions of 25 °C temperature, 24-h light and 80–90% humidity.

The first population, DB_4_, is a part of four population blocks (i.e. DB_1–4_) that trace their ancestry to the IV populations that were wild caught in Amherst, Massachusetts, USA (Ives 1970). The maintenance regime of the DB populations is available elsewhere (Sah, Salve & Dey 2013; Mishra *et al.* 2018b), and is mentioned here in brief. These populations (including DB_4_) are maintained under a 21-day discrete generation cycle, where 60–80 eggs are collected from the adult population cage of previous generation into 40 transparent 35-mL vials with ~6 mL of banana-jaggery medium. On the 12^th^ day after egg collection, the adults are transferred to a fresh plexi-glass cage that contains ~70 mL banana-jaggery medium in a 100-mm petri plate. These food plates are replaced with a fresh food plate every alternate day for the next six days, i.e. until the 18^th^ day since egg collection. On the 18^th^ day, the usual banana-jaggery food plate is supplemented with live yeast to boost the fecundity of female flies. On the 20^th^ day, the flies are provided with a food plate for oviposition over a window of ~14 hours. These eggs are then randomly sampled and distributed over 40 vials (as mentioned above) to start the next generation of the corresponding population.

The second population, named DBS [DB Scarlet-eyed], comprises individuals with a scarlet eye-color mutation. These flies were obtained from the ancestors of the DB populations that had been pooled together. At the time of this study, he DBS population diverged from the DB population for approximately 100 generations (~6 years). The maintenance of DBS population is identical to that of the DB populations, as described above.

The other two populations, VB_4_ and VBC_4_, have been derived from DB_4_ as a part of an ongoing dispersal-selection experiment. VB_4_ undergoes selection for higher dispersal every generation, while VBC_4_ serves as its corresponding control (Tung *et al.* 2018b). As a result of this selection, VB flies have evolved a higher dispersal propensity (see section 2.5 below) and locomotor activity than the corresponding VBC flies (Tung *et al.* 2018a; Tung *et al.* 2018b).

These populations have been previously used for studying context-dependent dispersal. For instance, it has been shown that density dependence and sex bias in dispersal is affected by the pre-dispersal context and presence of mates (Mishra *et al.* 2018b). Similarly, it is known that the difference in the dispersal of VB and VBC populations is affected by the presence or absence of resources (food and water) (Tung *et al.* 2018b).

### 2.2 Dispersal setup and assay

We used two-patch *source-path-destination* setups (*sensu* Mishra *et al.* 2018a; Mishra *et al.* 2018b; Tung *et al.* 2018b) to study fly dispersal. A *source* container (100-mL conical glass flask) is connected to a 2-m long *path* (transparent plastic tube; inner diameter ~1 cm), the other end of which opens into a *destination* container (250-mL plastic bottle) (Fig. S1; Mishra *et al.* 2018b). For the dispersal assays in this study, we introduced flies into the *source* and allowed them to disperse for 2 h. During this period, we replaced the *destination* container with a fresh, empty container every 15 min, with minimum possible disturbance to the rest of the setup. The number, sex and eye color (wherever applicable) of successful dispersers (flies found in the *destination* container) at each of these 15-min intervals were manually recorded. These data allowed us to compute the dispersal propensity and temporal profile of dispersers (see section 2.5 for more details).

This dispersal setup is based on extensive standardizations in the lab. Typically, the flies used in an experiment have *ad libitum* access to food and water (in the form of agar-based banana-jaggery medium) prior to the dispersal assay. Therefore, introducing them into an empty *source* container represents a change in the resource state of their environment. This mimics a situation where the local environment turns unfavorable or uninhabitable for a population, thereby promoting emigration from the *source* container.

### 2.3 Rearing flies for the experiments

To eliminate any confounding effects of habitat quality, the entire dispersal setup (section 2.2) was devoid of food and moisture, and the identity of individuals in the source was the only difference across treatments during the dispersal assay. As dispersal is known to be affected by factors such as age (Hastings 1992), density (Matthysen 2005), kin competition (Gandon 1999) and level of inbreeding (Charlesworth & Charlesworth 1987), we avoided these potential confounds through appropriate rearing of the flies (detailed in Supplementary Text S1).

### 2.4 Experimental design

Two experiments were performed to investigate the behavioral plasticity of male and female flies for dispersal. In Experiments 1 and 2, we examined the change in male and female dispersal under mixed-sex and same-sex conditions, respectively.

#### 2.4.1 Experiment 1: Mixed-sex dispersal

This experiment aimed to discern how dispersal capability of one sex affects the dispersal of the other sex through differential mate-availability. Dispersal of males from the baseline population (DB_4_) was measured in the presence of either dispersal-selected females (VB_4_) or control females (VBC_4_) that were not selected for dispersal. Similarly, dispersal of baseline females (DB_4_) was examined in the presence of either more-dispersive (VB_4_) or less-dispersive (VBC_4_) males. Thus, this experiment had four treatments: [VB F+ DB M] and [VBC F + DB M], and [VB M + DB F] and [VBC M + DB F], where M stands for males and F denotes females (Fig. 1). The initial sex ratio in the source was 1:1 for each treatment, comprising 60 males and 60 females. Following an earlier protocol (Mishra *et al.* 2018b), we carried out the experiment over multiple consecutive days, with a fresh set of age-matched adult flies (12-day-old from the date of egg collection) every day. Twenty-one hours prior to the dispersal assay on each day, a fresh batch of newly eclosed flies was separated by sex under light CO_2_ anesthesia and maintained in same-sex groups of 60 individuals. Right before the dispersal assay, 60 males and 60 females from the relevant populations were mixed to yield the corresponding treatments. The experiment ran for seven consecutive days, with one replicate/treatment on each day, resulting in seven independent replicates for each treatment. In total, 3,360 flies (2 sexes × 4 treatments × 7 replicates × 60 flies sex^−1^ treatment^−1^ replicate^−1^) were used for this experiment.

**Fig. 1.**
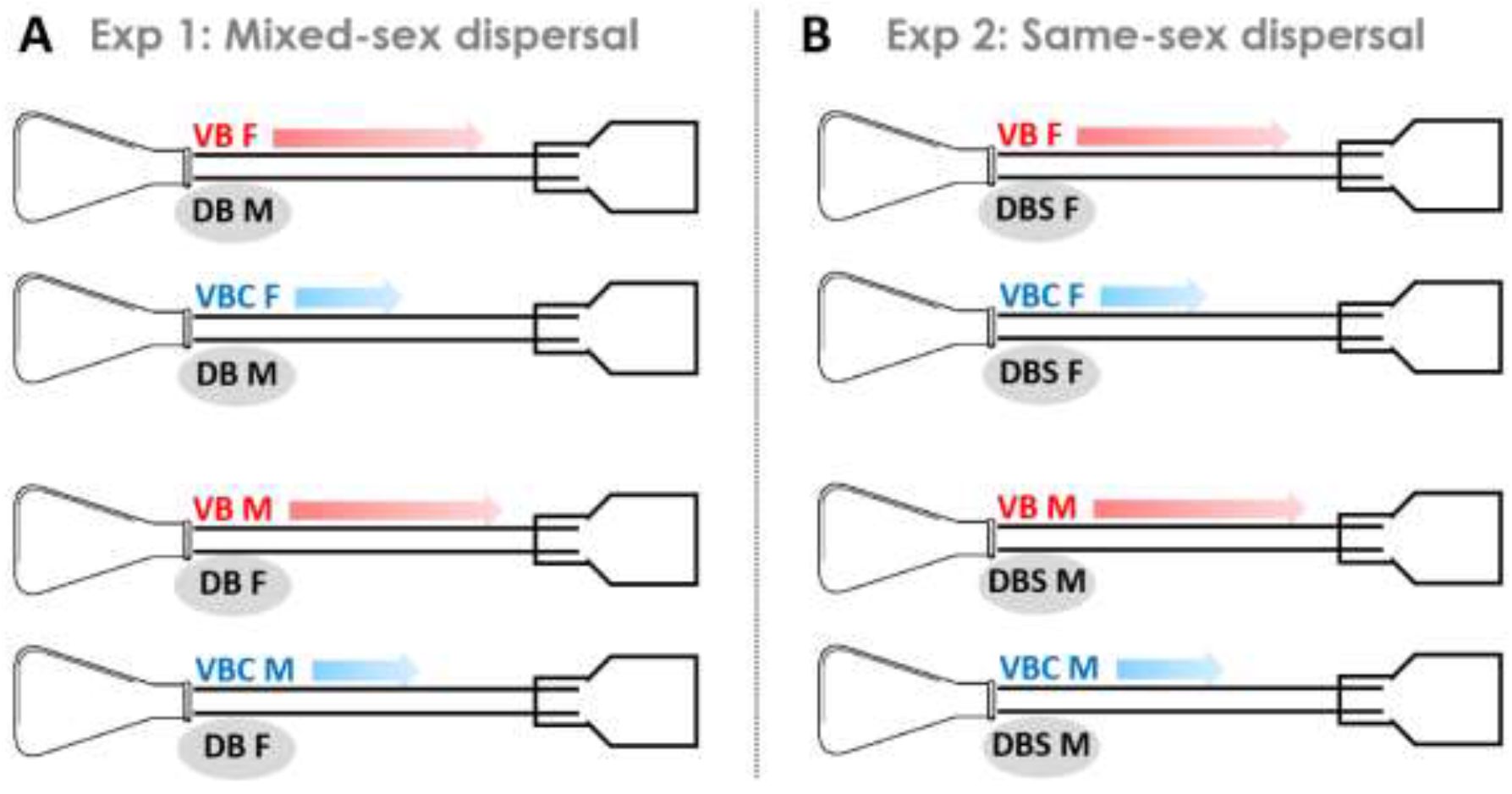
Schematics of the experimental design. (A) In Experiment 1, we studied how the dispersal of one sex was affected by the dispersal of the other sex. The dispersal of DB flies (baseline population) was investigated in the presence of VB (dispersal-selected) vs. VBC (control) flies of the opposite sex. The arrow lengths denote the high-dispersive vs. low-dispersive nature of the VB vs. VBC individuals. (B) In Experiment 2, the behavioral plasticity of dispersal was tested in the context of differential dispersal by same-sex individuals. The dispersal of DBS (Scarlet-eyed baseline population) was investigated in the presence of VB (dispersal-selected) vs. VBC (control) flies of the same sex.

#### 2.4.2 Experiment 2: Same-sex dispersal

In this experiment, we investigated whether dispersal of a sex (male or female) was affected by the dispersal of other individuals of the same sex. Similar to Experiment 1 (section 2.4.1), we compared the dispersal of baseline flies in the presence of dispersal-selected vs. control flies. However, since this experiment would comprise individuals of only one sex in each treatment, we used flies with a scarlet eye-color mutation (population DBS) instead of DB_4_, to distinguish between the baseline flies and VB or VBC flies. The rest of the protocol was identical to the one followed in Experiment 1, and the four treatments in this case were: [VB F + DBS F] and [VBC F + DBS F], and [VB M + DB M] and [VBC M + DB M] (Fig. 1). As in Experiment 1, we used 3,360 flies (2 eye-colors × 4 treatments × 7 replicates × 60 flies eye color^−1^ treatment^−1^ replicate^−1^) for this experiment.

### 2.5 Dispersal traits

For every dispersal setup (replicate), we recorded the number and the sex (Experiment 1) or eye color (Experiment 2) of flies that reached the *destination* during each 15-min interval of the dispersal assay (lasting 2 h). In addition, we also recorded these data for flies that emigrated from the source but did not reach the destination, i.e. the flies found within the *path* at the end of dispersal assay.

We then estimated the dispersal propensity, i.e. the proportion of flies that initiated dispersal (i.e. emigrated) from the source (Friedenberg 2003), as:

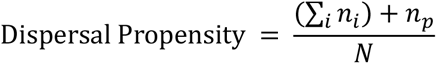

where *n*_*i*_ is the number of flies that reached the destination during the *i*^th^ 15-min interval, *n*_*p*_ is the number of flies found within the path at the end of dispersal assay and *N* is the total number of flies introduced in the setup (here, 120). Therefore, the dispersal propensity can also be considered as the emigration probability from the source, a term that has been used in the literature in the past (e.g. Englund & Hambäck 2004). However, here we prefer to use dispersal propensity to refer to this property.

In addition, we also estimated the overall temporal distribution of dispersers reaching the destination (similar to Mishra *et al.* 2018b). For this, the number of dispersers that reached the destination during each 15-min interval was divided by the final number of successful dispersers. Finally, following a suggestion by a reviewer, we also estimated the dispersers in each time bin as a proportion of remaining individuals (i.e. those who had not reached the destination until that point).

### 2.6 Statistical analyses

As stated above, both Experiment 1 and Experiment 2 were performed over seven consecutive days, with one replicate of each treatment assayed every day. Therefore, we analyzed these data with *replicate* as a random blocking factor. For each experiment, the dispersal propensity of VB and VBC flies were first compared in a single analysis, to establish the difference in biotic environment faced by DB (or DBS) flies while dispersing with VB or VBC individuals. The dispersal propensity data for DB (or DBS) flies were then compared in another analysis, to assess mate-finding dispersal (or effect of same-sex dispersal).

We fit Generalized Linear Mixed Models (GLMMs) with binomial error distribution (and logit link function) for the dispersal propensity data, using the ‘glmmPQL’ function in the ‘MASS’ package v7.3-51.4 (Ripley *et al.* 2013) in R v3.6.2 (R Core Team 2019). For a given comparison (e.g. VB/VBC) within a given experiment, we started with a full model, and then carried out marginality-based model reduction (following Crawley 2012; Pekár & Brabec 2016) until a minimal adequate model (MAM) was obtained.

In Experiment 1, the analysis for VB/VBC comparison had *sex* (male/female) and *dispersal selection* (VB/VBC) as the fixed factors, and *replicate* (1–7) as a random factor. Similarly, the analysis for DB individuals had *sex* (male/female) and *mate dispersal* (VB/VBC) as the fixed factors, and *replicate* (1–7) as the random factor. The corresponding analyses for Experiment 2 were similar to those of Experiment 1, except for one difference. As each treatment in Experiment 2 comprised only one sex, the analysis for DBS individuals had *same-sex dispersal* (VB/VBC) as a fixed factor instead of *mate dispersal* (VB/VBC). The other fixed factor (*sex*) and the random factor (*replicate*) remained the same.

For analyzing the temporal profiles of dispersers, we again used binomial GLMMs, similar to the framework described above for dispersal propensity data. However, in addition to the relevant fixed and random factors for each case, *time* (15, 30, 45, 60, 75, 90, 105, and 120 min) was used as an additional fixed factor in all the analyses. Finally, similar to the protocol followed above, marginality-based model reduction was carried out in each case, until an MAM was obtained.

Cohen’s *d* was used as a measure of effect size for pairs of groups, and the effect was interpreted as large, medium and small for *d* ≥ 0.8, 0.8 > *d* ≥ 0.5 and *d* < 0.5, respectively (Cohen 1988).

## 3. Results

### 3.1 Significant behavioral plasticity in dispersal for both sexes in the mixed-sex experiment

In Experiment 1, we investigated how the dispersal of males or females was affected by the movement of the opposite sex. For this, we first established the difference between the dispersal of VB (dispersal-selected) and VBC (control) individuals, to assess the extent of difference experienced by DB (baseline) individuals. This was followed by an analysis of DB dispersal across the four treatments, to examine the effect of dispersing with high-vs. low-dispersive mates. We report the results from Minimal Adequate GLMM fits here in the main text, while the Supplementary Information contains the details of model reduction (Text S2 and S3).

As expected from previous studies (Tung *et al.* 2018a; Tung *et al.* 2018b), the Minimum Adequate Model (MAM) for dispersal propensity revealed a significant main effect of *dispersal selection* (t = −5.99, *P* < 10^−5^), with VB individuals being more dispersive than VBC individuals. The effects of both *sex* and *dispersal selection × sex* interaction were found to be non-significant at various stages of model selection (Text S2.1), thereby excluding them from the MAM. This suggests that the effect of dispersal selection was symmetric across males and females (Figs 2A and 2B). The difference between dispersal-selected and control flies was also apparent in their temporal dispersal profiles, as majority of VB dispersers completed the dispersal much faster than the VBC dispersers (Text S3.1) (Figs 3A and 3B; see Supplementary Figs S2A and S2B for a different representation of the same data). These differences meant that the DB flies would have faced markedly different temporal mate availability, depending on whether they were dispersing with VB or VBC individuals of the opposite sex. The dispersal data of DB individuals were next analyzed to capture the effects of this contrast in biotic environments.

**Fig. 2.**
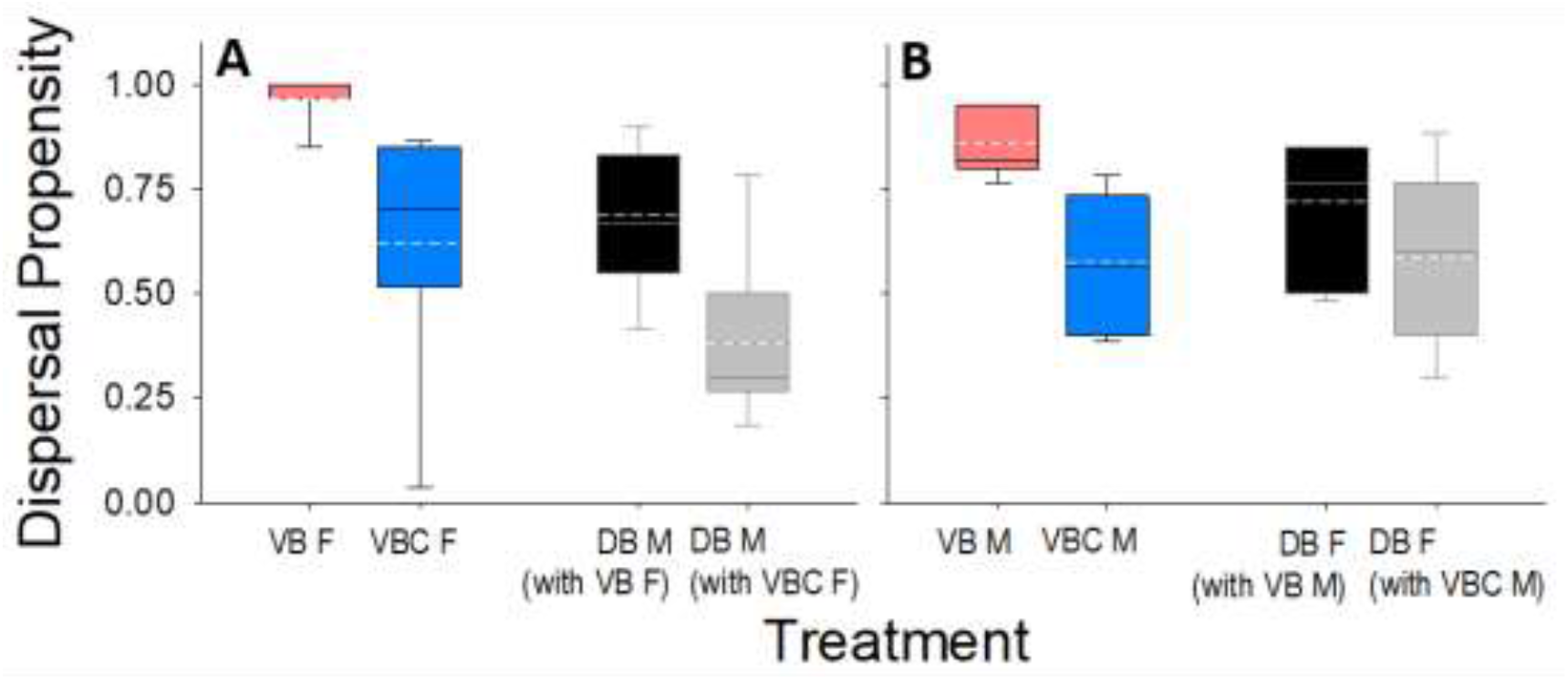
Significant behavioral plasticity observed in mixed-sex dispersal (Experiment 1). Box-plots for the dispersal propensity of (A) VB (high-dispersive) vs. VBC (low-dispersive) females, and that of the DB (baseline) males dispersing with them, (B) VB (high-dispersive) vs. VBC (low-dispersive) males, and that of the DB (baseline) females dispersing with them. The edges of the box in each case denote the 25^th^ and 75^th^percentiles, while the solid line and dashed line represent the median and the mean, respectively.

**Fig. 3.**
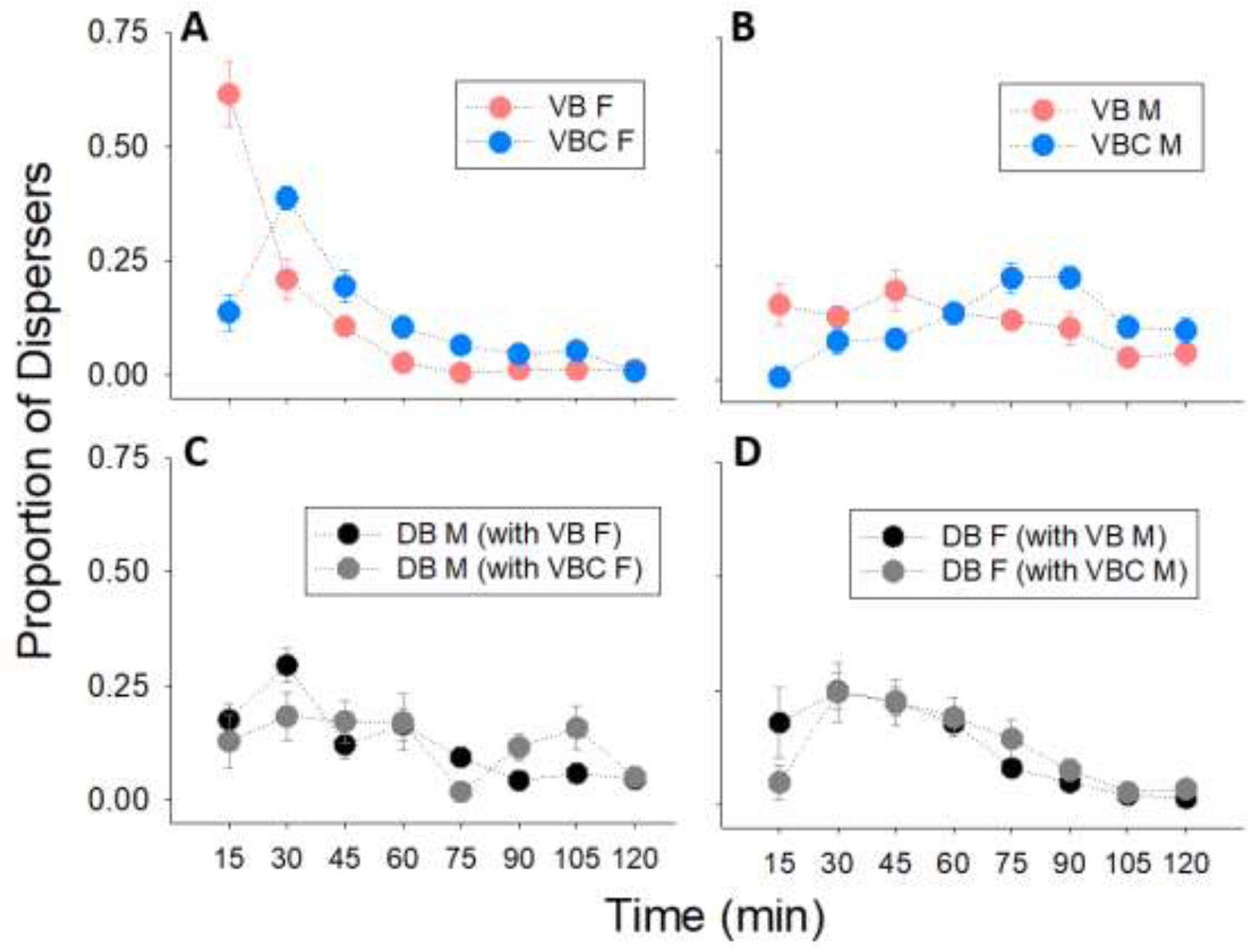
Temporal dispersal profile in the mixed-sex scenario (Experiment 1). Proportion of individuals reaching the destination over the 2-h dispersal period, for (A) VB (high-dispersive) vs. VBC (low-dispersive) females, (B) VB (high-dispersive) vs. VBC (low-dispersive) males, (C) DB (baseline) males dispersing with VB vs. VBC females, and (D) DB (baseline) females dispersing with VB vs. VBC males. See Supplementary Text S3 for the associated analyses.

The MAM for DB individuals revealed a significant effect of *mate identity* (t = −4.18, *P =* 0.0005), as their dispersal propensity in the presence of VB mates was much higher than that with VBC mates (Figs 2A and 2B). There was a significant main effect of *sex* as well (t = −2.34, *P =* 0.030), with females showing higher overall dispersal than males. The MAM excluded the *mate identity × sex* interaction, as its effect was not significant in the full model (Text S2.2). However, the effect size of the increase in DB dispersal was larger in males than in females (males: d=1.73 (large); females: d=0.76 (medium)) (Figs 2A and 2B). This was also reflected in the temporal dispersal profiles of the DB individuals, where DB males showed a greater change in the timing of their dispersal than DB females (Text S3.2) (Figs 3C and 3D; see Supplementary Figs S2C and S2D for a different representation of the same data). Thus, dispersal was found to be plastic in both sexes, as assessed in the presence of high-vs. low-dispersive mates, although the magnitude of this effect was larger in males.

### 3.2 No significant behavioral plasticity in dispersal for either sex in the same-sex experiment

Similar to Experiment 1, the dispersal of VB and VBC individuals was first compared across the four treatments in Experiment 2. Thereafter, data for DBS [DB Scarlet-eyed] individuals were analyzed, to test if their dispersal was affected by high-vs. low-dispersive individuals of the same sex.

As observed in Experiment 1, the MAM revealed a significant main effect of *dispersal selection* (t = −4.56, *P =* 0.0002), as the dispersal propensity of VB individuals was higher than that of VBC individuals. There was also a significant effect of *sex* (t = −2.18, *P =* 0.042), as the overall dispersal propensity was higher in females than in males. Finally, the *dispersal selection × sex* interaction was not significant in the full model, and therefore, excluded from the MAM (Text S2.3; Figs 4A and 4B). However, difference in the temporal profile of VB and VBC dispersers was apparent only in males and not in females (Text S3.3) (Figs 5A and 5B; see Supplementary Figs S3A and S3B for a different representation of the same data).

**Fig. 4.**
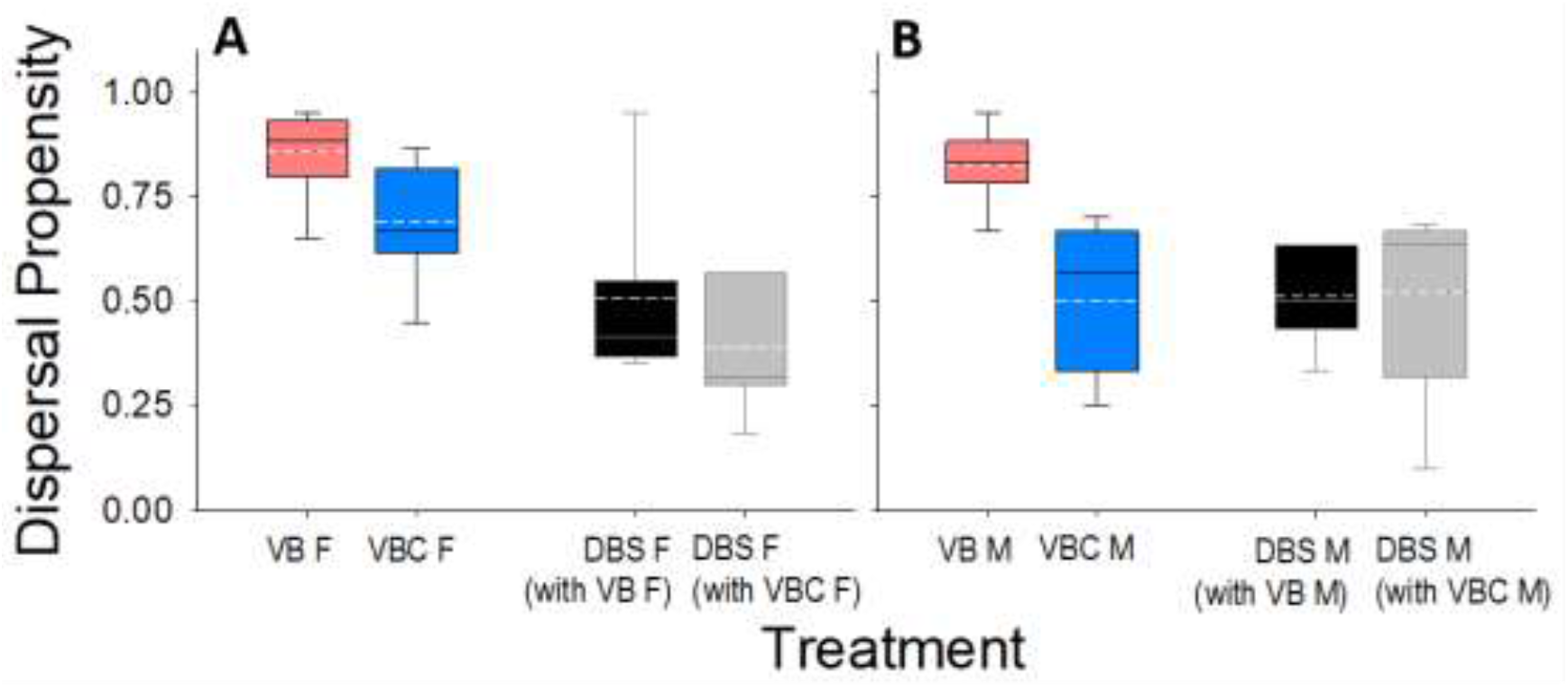
No significant behavioral plasticity in same-sex dispersal (Experiment 2). Box-plots for the dispersal propensity of (A) VB (high-dispersive) vs. VBC (low-dispersive) females, and that of the DBS (scarlet-eyed baseline) females dispersing with them, (B) VB (high-dispersive) vs. VBC (low-dispersive) males, and that of the DBS (scarlet-eyed baseline) males dispersing with them. The edges of the box in each case denote the 25^th^ and 75^th^percentiles, while the solid line and dashed line represent the median and the mean, respectively.

**Fig. 5.**
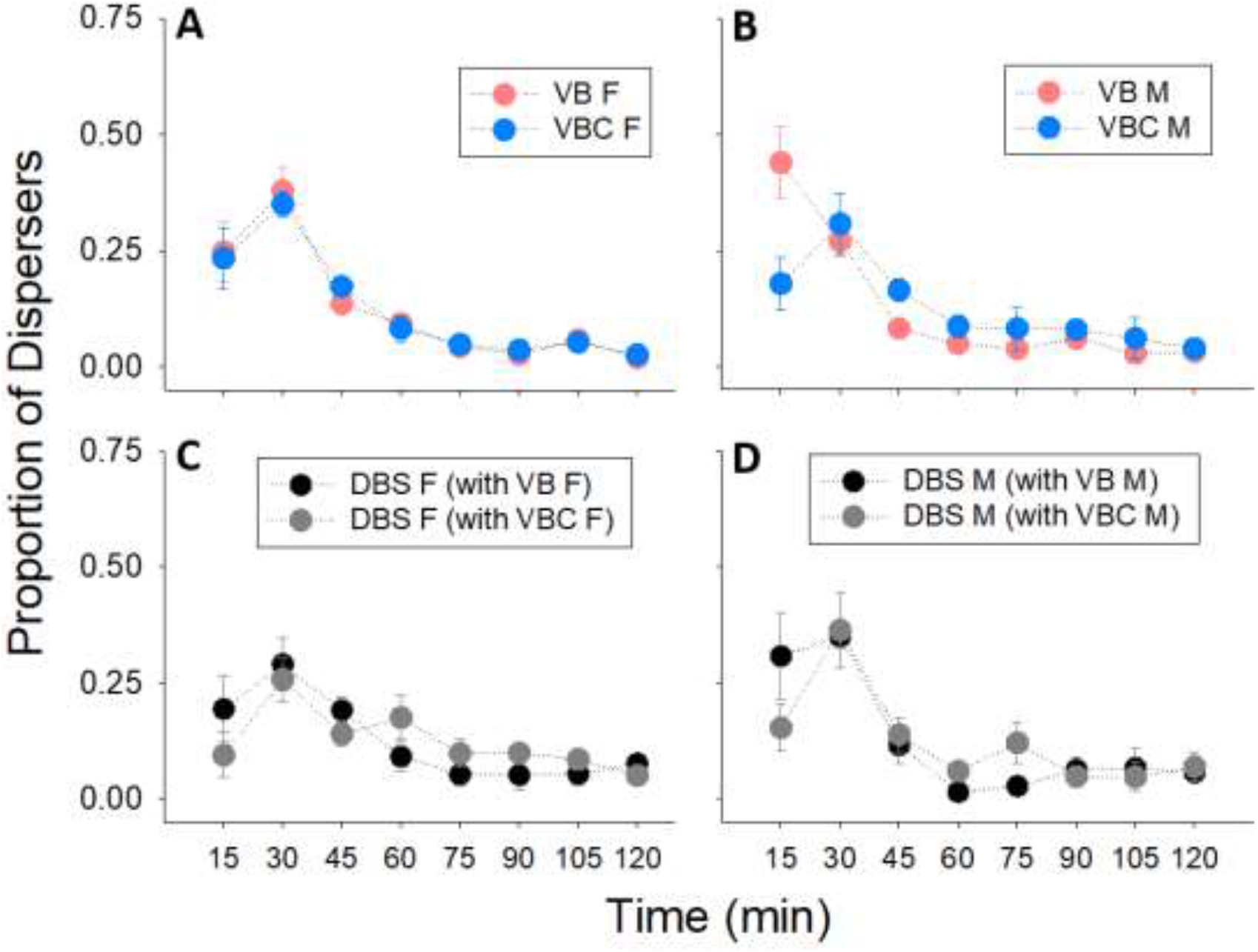
Temporal dispersal profile in the same-sex scenario (Experiment 2). Proportion of individuals reaching the destination over the 2-h dispersal period, for (A) VB (high-dispersive) vs. VBC (low-dispersive) females, (B) VB (high-dispersive) vs. VBC (low-dispersive) males, (C) DBS (scarlet-eyed baseline) females dispersing with VB vs. VBC females, and (D) DBS (scarlet-eyed baseline) males dispersing with VB vs. VBC males. See Supplementary Text S3 for the associated analyses.

For DBS flies, neither of the three effects, i.e. *same-sex dispersal*, *sex*, and *same-sex dispersal × sex*, were significant at any stage of model selection (Text S2.4). The final MAM had included only a non-significant effect of *sex* (t = 1.01, *P* = 0.32). This implied that dispersal of either sex was not affected by high-vs. low-dispersive individuals of the same sex. This is despite the fact that the temporal profile of DBS males and females registered some difference across the treatments (Text S3.4) (Figs 5C and 5D; see Supplementary Figs S3C and S3D for a different representation of the same data).

## 4. Discussion

### 4.1 Mate-finding dispersal observed in the presence of differential mate dispersal

The primary result of our study is that mate dispersal (and consequently, local mate availability) is a key proximate determinant of dispersal. In Experiment 1, we studied the dispersal of males and females of the baseline populations (DB) in the presence of high-dispersive (VB) and low-dispersive (VBC) mates. DB individuals showed significantly higher dispersal in the presence of high-dispersive mates than in the presence of low-dispersive mates (*cf* Figs 2 and 4). While these results from Experiment 1 support the hypothesis that dispersal is influenced by different levels of mate dispersal, they do not constitute sufficient evidence for the same. This is because these results could simply be a manifestation of negative density-dependent dispersal, wherein per-capita dispersal increases with decreasing density (e.g. Baguette, Clobert & Schtickzelle 2011; Mishra *et al.* 2018b). In such a scenario, the observations could be explained without invoking the role of mates. This possibility was ruled out by Experiment 2, where mate availability was not a factor and we could test the effect of same-sex individuals on dispersal. In contrast to the results of Experiment 1, Experiment 2 showed that neither males nor females showed a difference in dispersal in the presence of same-sex individuals with different dispersal levels. Thus, taking together the results from Experiments 1 and 2, we concluded that mate dispersal, and not the dispersal of same-sex individuals, modulates dispersal in *D. melanogaster*. We do note a minor caveat here – although the DBS population shares its ancestry and maintenance regime with the DB population (Section 2.1), the difference in the results of the two experiments could, in principle be in part due to some population-specific differences.

While we used populations with differential dispersal propensity (i.e. VB and VBC) to achieve Sex-Biased Dispersal (SBD) and create differences in local mate availability, a few previous studies have tried using treatments with skewed sex ratios for the same effect. For instance, Trochet *et al.* (2013) used three different sex-ratio treatments to study dispersal in the butterfly, *Pieris brassicae*. Similarly, Baines *et al.* (2017) used treatments with equal, female-biased and male-biased sex-ratios to assess changes in SBD of a semiaquatic insect, *Notonecta undulata*. However, in both these studies, the authors reported that sex ratio treatments were not effective in producing or modulating SBD. The authors then hypothesized that other mechanisms, including competition and dispersal costs, were likely more important reasons for dispersal than local mate availability (Trochet *et al.* 2013; Baines, Ferzoco & McCauley 2017). Since in our study, none of the treatments started with a skewed ratio, and rather, mate availability changed through time (as evidenced by the temporal dispersal profiles in Figs 3A and 3B), it is possible that active “mate-following” led to mate-finding dispersal.

As mate limitation is believed to be the most common cause of Allee effects in sexually reproducing species (Taylor & Hastings 2005; Berec *et al.* 2018), several theoretical studies have investigated the effects of mate limitation on population growth and spread, as well as the evolution of mate-finding dispersal. For instance, mate limitation and its effects are predicted to be especially pronounced in species with monogamous mating systems (Shaw & Kokko 2014; Shaw, Kokko & Neubert 2018). However, we observed these effects in our *D. melanogaster* populations, which are not only polygamous, but their rearing and age was such that they had most likely mated by the time of the experiment (Supplementary Text S1). As a result, the mate limitation faced by them was likely not very severe. Furthermore, adaptations other than mate finding, such as multiple matings, long mating window and sperm storage, are expected to mitigate mate limitation (Gascoigne *et al.* 2009; Shaw & Kokko 2014; Fromhage, Jennions & Kokko 2016). All these traits are well documented in *Drosophila* (Fuerst, Pendlebury & Kidwell 1973; Pyle & Gromko 1978; Pitnick, Marrow & Spicer 1999). Finally, this also means that the relative order of mating and dispersal, which is a central basis for many theoretical studies on mate-finding and sex-biased dispersal (Shaw & Kokko 2014; Li & Kokko 2019), may not be a very crucial factor for *Drosophila*, which have a long mating window and can potentially mate before, during or after dispersal. Therefore, mate-finding dispersal could be even more significant in those taxa where some or all of these traits are less pronounced (Fromhage, Jennions & Kokko 2016).

### 4.2 Sex differences in mate-finding dispersal

Once mate-finding dispersal had been established, the next objective was to see if this behavior was symmetric between males and females. Our results from Experiment 1 showed that the effect size of mate-finding dispersal was much larger for males than for females (section 3.1, *cf* Fig. 2A and Fig. 2B). A straightforward explanation for this observation could be the differences in the temporal dispersal profile of VB vs. VBC males and females. As can be seen in Figs 3A and 3B, more than 60% dispersers among the VB females completed dispersal within the first 15 minutes. Coupled with the high dispersal propensity of VB females (Fig. 2A), this means that within the first few minutes of the experiment, the sex ratio in the *source* of [VB F + DB M] treatment deviated significantly from 1:1, leading to the DB males in this treatment facing a shortage of females. In contrast, it took 45 min for ~60% of the dispersers among the VB males to complete dispersal (Fig. 3B). As a result, the extent of mate shortage created by the dispersal of VB males was likely not as severe as that created by the dispersal of VB females. Another possible reason for the greater mate-finding dispersal observed in DB males is that dispersal propensity of DB females was inherently higher than that of DB males. This can be observed by comparing the DB male data in the presence of low-dispersive (VBC) females in Fig. 2A with the DB female data in the presence of low-dispersive (VBC) males (Fig. 2B). As a result, the available scope for increase in dispersal was likely limited for females compared with males. This is possible because, in our setup, males were dispersing only to escape the desiccation and starvation stress, whereas females had a dual rationale for dispersal: escape from stress as well as the search for a suitable oviposition site. Finally, since our flies were likely already mated at least once, the males probably had more to gain from any extra matings than the females. In fact, extra matings could even lead to deleterious physiological consequences for the females, as evidenced by the extensive male mate-harm literature in *Drosophila* (Levine *et al.* 1980; Harshman, Hoffmann & Prout 1988; Kuijper, Stewart & Rice 2006). As a result, the higher inherent dispersal by DB females might also be a way to escape from excessive male mate harm (e.g. see Byrne, Rice & Rice 2008). Overall, these results are in line with the expectations of “male-biased mate searching” from earlier theoretical studies, which posit that females disperse primarily for resources, whereas males disperse primarily for mates (Meier, Starrfelt & Kokko 2011; Hovestadt, Mitesser & Poethke 2014; Fromhage, Jennions & Kokko 2016).

Our current results are also relevant in the context of dispersal evolution. We had earlier reported that SBD did not evolve in the VB populations, even though a two-patch *source-path-destination* setup like ours could potentially select for female-biased dispersal (Tung *et al.* 2018b). This is because, in principle, the males could maximize their fitness by mating with as many females as possible before the dispersal run and avoid the dispersal costs altogether, whereas the females had no recourse but to complete the dispersal from *source* to *destination* to realize their fitness (Shaw & Kokko 2014; Tung *et al.* 2018b). Our current results demonstrate how strong mate-finding dispersal would have countered this asymmetry in the selection pressure between males and females, thus, hindering the evolution of SBD. This is also in line with the theoretical predictions that the two sexes should evolve similar dispersal kernels over time (Meier, Starrfelt & Kokko 2011; Shaw & Kokko 2014). Therefore, our current results highlight the role of mate-finding dispersal in modulating phenomena such as the evolution of SBD.

### 4.3 Implications of our results

Our results revealed a strong effect of mate dispersal, more so in males than in females. To our knowledge, this is the first clear demonstration of how mate-finding dispersal, can counter the sex bias in dispersal. We discuss some implications of our results below.

First, SBD is known to be a major cause of differential mate availability (Meier, Starrfelt & Kokko 2011; Miller *et al.* 2011; Miller & Inouye 2013). Here we show the effects of SBD on mate availability can be countered over short time scale via mate-finding dispersal. Miller & Inouye (2013) hypothesized that demographic stochasticity can be a major factor that dampens the effects of SBD on mate availability. We show that a more deterministic factor could be demographic rescue via mate-finding dispersal. The extent of such dispersal-mediated demographic rescue, in turn, would depend on the degree of patch isolation and costs of dispersal (Gascoigne *et al.* 2009).

Second, mate limitation is especially important for small populations, particularly those found at invasion fronts and range boundaries. SBD can lead to skewed sex ratios at invasion fronts (Miller & Inouye 2013), with acute mate shortage resulting in strong Allee effects (Taylor & Hastings 2005; Contarini *et al.* 2009). As a result, it has been suggested that SBD makes invasions more variable (Miller & Inouye 2013). However, our results indicate that such variation would be limited in the presence of deterministic factors such as mate-finding dispersal. In fact, mate-search efficiency is predicted as a key parameter that determines population-level effects including growth and spatial spread (Shaw, Kokko & Neubert 2018). Overall, we hypothesize that the effect of mate-finding dispersal on invasions would depend on population size. While mate-finding dispersal can rescue small populations from Allee effects, it might dilute the process of dispersal evolution via spatial sorting (Shine, Brown & Phillips 2011) in larger populations. This is because mate-finding dispersal is “context-dependent”, as opposed to “phenotype-dependent” (*sensu* Clobert *et al.* 2009; Clobert 2012), which would imply a low heritability of dispersal traits among the dispersers. If spatial sorting is diluted in this manner, it is expected to slow down the speed of expansions.

Third, sex differences have been reported in life-history/behavioral traits related to dispersal (Legrand *et al.* 2016; Mishra *et al.* 2018a). Sex bias in mate-finding dispersal can thus interact with these sex differences in life history or behavior, to modulate the final distribution of individuals in spatially structured populations. *Inter alia*, such interactions can amplify/weaken the Allee effects experienced by the population (Shaw & Kokko 2014). Thus, elucidating the mechanisms of sex bias in mate-finding dispersal (e.g. Shaw & Kokko 2014; Fromhage, Jennions & Kokko 2016) would be an important factor in understanding the nature of these interactions and the consequent population-level effects.

Finally, it should be noted that results from microcosm experiments such as ours should not be extrapolated directly to natural populations of even the same taxon. Natural populations often face a number of environmental conditions at once: for instance, in nature, factors such as wind, odour cues and predators may play a role in shaping *Drosophila* dispersal (Dobzhansky 1973; Dickinson 2014). That is why observational studies in the field need to adopt a “top-down” approach, where the system data are decomposed into possible relevant components. In contrast, microcosm studies follow a “bottom-up” approach, i.e. simplifying the system as much as possible, and studying the effect of very few factors (typically, one or two) in detail. Therefore, extrapolating our results to natural populations of *Drosophila* was neither our objective nor within the scope of our current study. The strength of our experiments lies in the fact that they allow us to remove confounding factors such as passive dispersal (e.g. by wind), extraneous cues (e.g. predators or other food sources), and effects of density, thereby allowing us to focus exclusively on the ecological issue of mate-finding dispersal.

## Acknowledgements

We thank Mohammed Aamir Sadiq and P. M. Shreenidhi for help with the experiments and two anonymous reviewers for helpful comments. ST and AM were supported by Senior Research Fellowships from the Council of Scientific and Industrial Research, Government of India. VRSS was supported through the GE Foundation Scholar Leaders Program. SS was supported through the INSPIRE fellowship of Department of Science and Technology (DST), Government of India. This study was supported by a research grant (#CRG/2018/001333) from Science and Engineering Research Board, DST, Government of India, and internal funding from IISER-Pune.

## Authors’ Contributions

AM and SD conceived the ideas and designed methodology; AM, ST, VRSS and SS collected the data; AM and SD analyzed the data; AM and SD led the writing of the manuscript. All authors contributed critically to the drafts and gave final approval for publication.

## Supplementary Information

### Text S1: Rearing conditions of the flies used in experiments

Two cages, with ~2400 adult flies each, were maintained for each of the populations used (DB_4_, DBS, VB_4_ and VBC_4_). To collect eggs for a particular day of dispersal assay, one cage of each population was supplied with live yeast plate for ~24 h. Fresh banana-jaggery medium was then supplied to the flies, allowing the females to oviposit for 12 h. After this period, ~50 eggs each were collected into forty 35-mL plastic vials containing ~6 mL of banana-jaggery medium. Following this, a plate with live yeast paste was provided to the flies again, to enable another round of egg collection. The same procedure was followed for the other cage of a population as well, with a constant difference of 24 h between the cages. Therefore, it was possible to collect eggs every day, alternatively from two cages of the same population, for 7 days. The large breeding population size (~2400) in each cage ensured that the flies were not inbred, and random sampling from a large number of eggs during the collection reduced the chances of kin being sampled together. Moreover, as flies from a single set of collected eggs were used for the dispersal assay on a particular day, they were all of the same age (12^th^ day from egg collection) and likely mated at least once by the time of the assay.

**Fig. S1.**
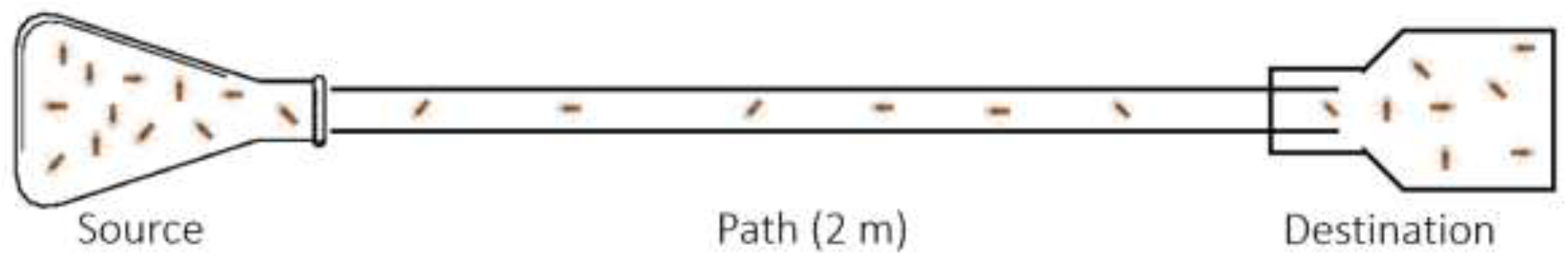
Two-patch dispersal setup used in the study. Adult individuals of *Drosophila melanogaster* are introduced in the *source* container (100-mL conical glass flask), the opening of which was connected to the *path* (transparent plastic tube of internal diameter ~1 cm). The other end of the *path* opened into the *destination* container (250-mL plastic bottle), with a protrusion of ~3 cm to prevent backflow of successful dispersers into the *path*. During the dispersal assay, the *destination* container could be replaced periodically with a fresh container, to estimate the temporal profile of successful dispersers. This schematic is similar to the one used in an earlier study (Mishra *et al.* 2018).

## Text S2: Model reduction details for dispersal propensity analyses

### Text S2.1: Experiment 1 (VB/VBC comparison)

#### 1) Full model

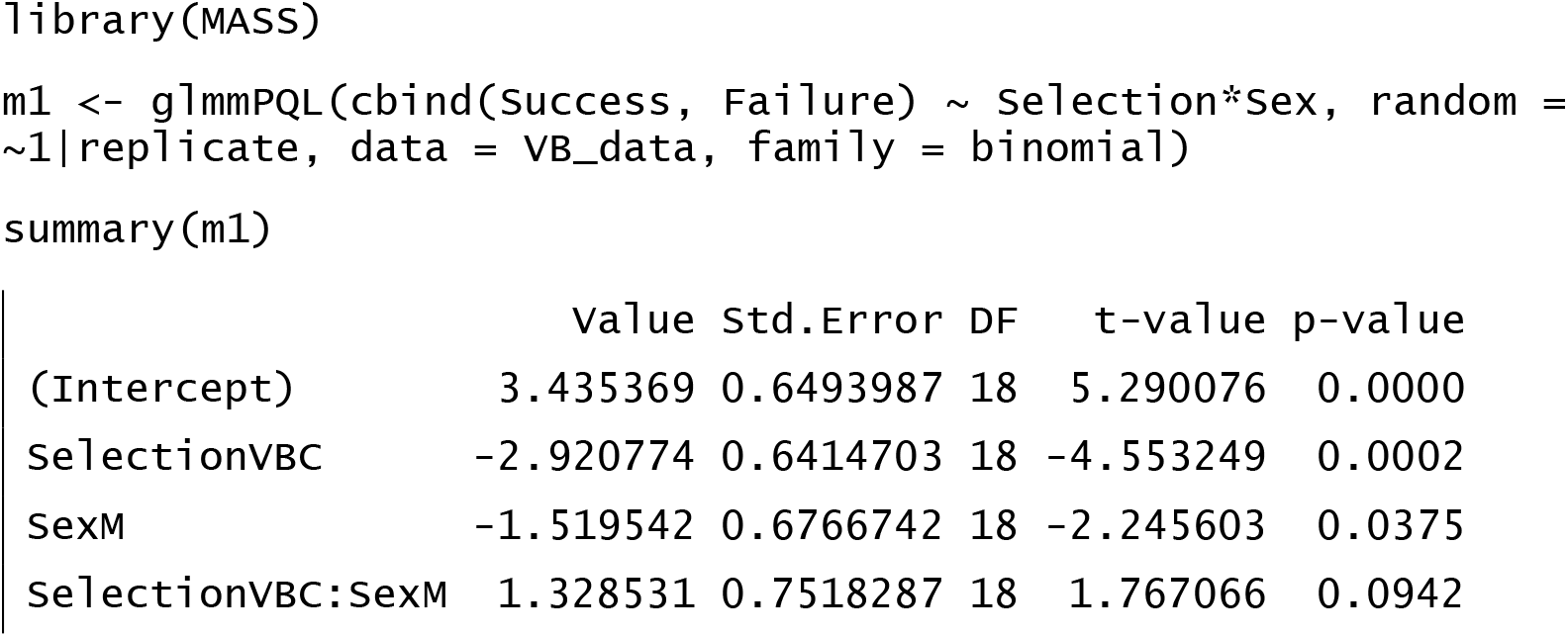

#### 2) Model reduction (removing the non-significant *dispersal selection × sex interaction*)

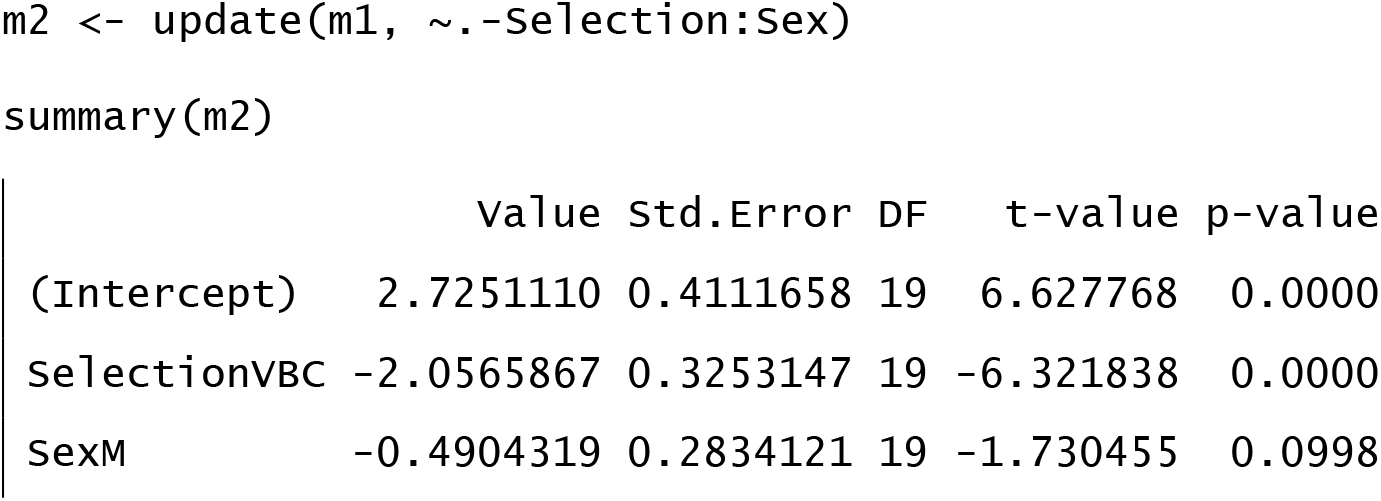

#### 3) Model reduction (removing the non-significant factor, *sex*)

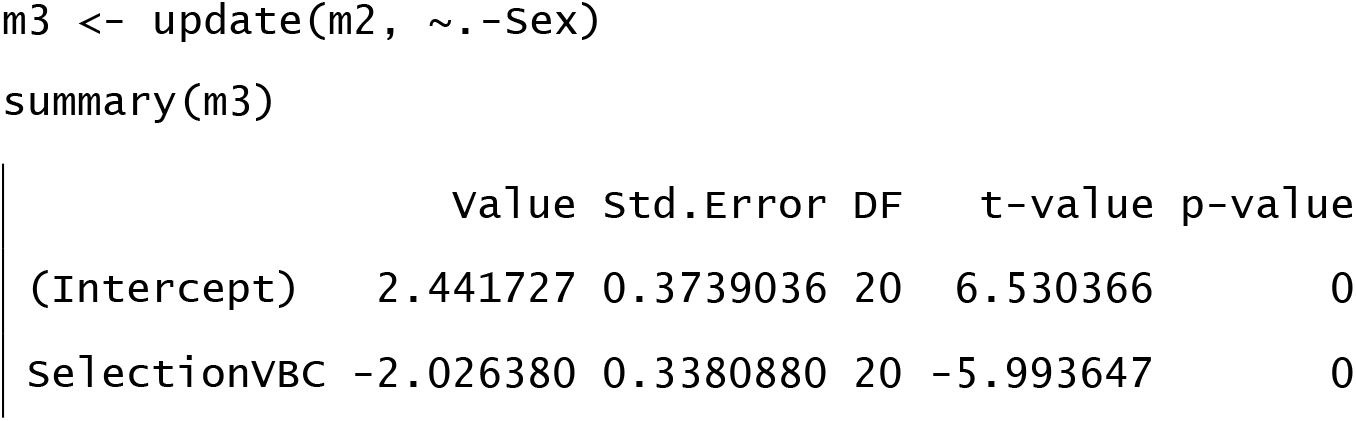

#### Minimum Adequate Model (MAM): m3

### Text S2.2: Experiment 1 (DB comparison)

#### 1) Full model

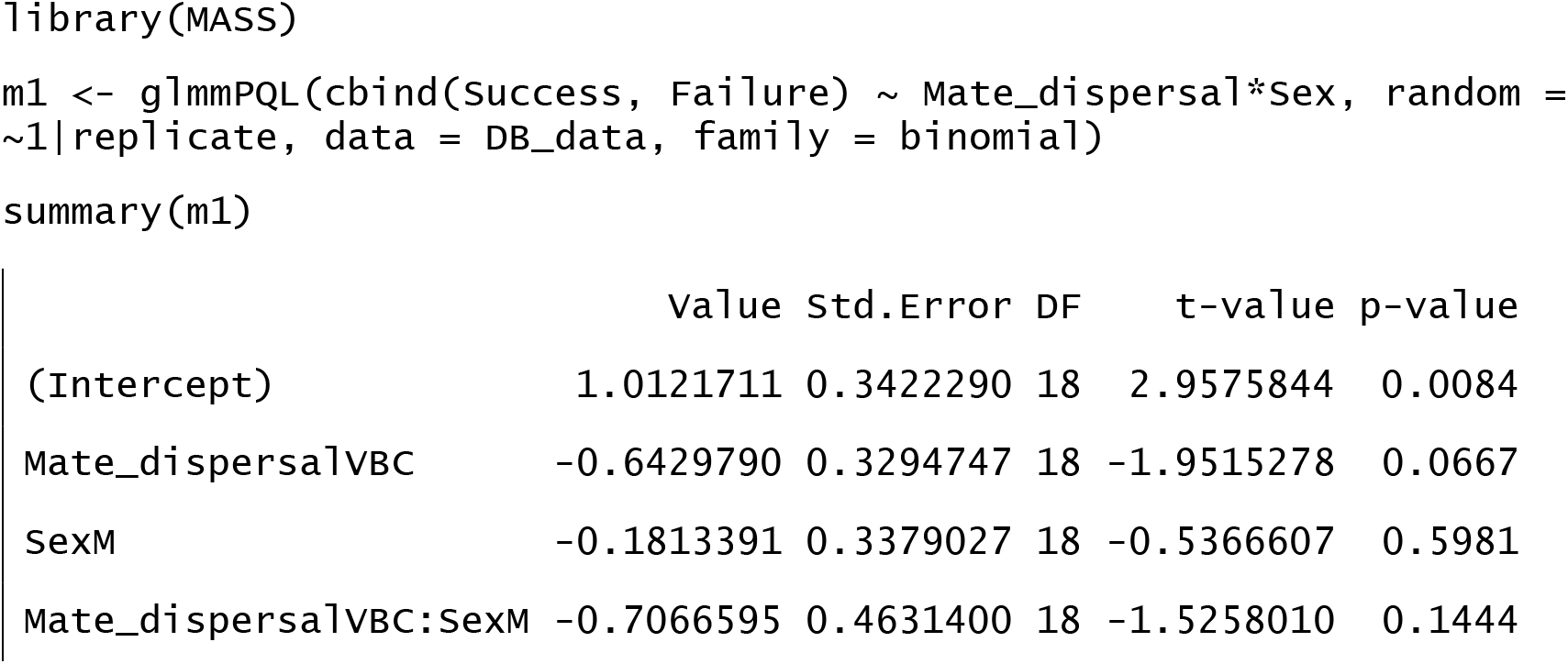

#### 2) Model reduction (removing the non-significant *mate dispersal × sex* interaction)

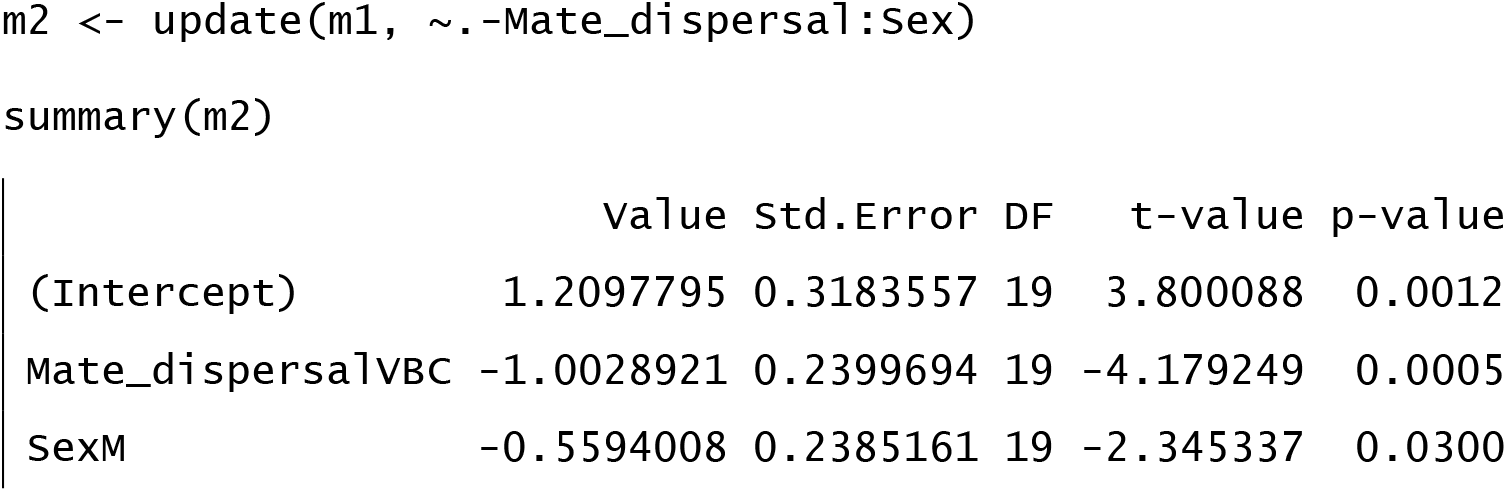

#### 3) Minimum Adequate Model (MAM): m2

### Text S2.3: Experiment 2 (VB/VBC comparison)

#### 1) Full model

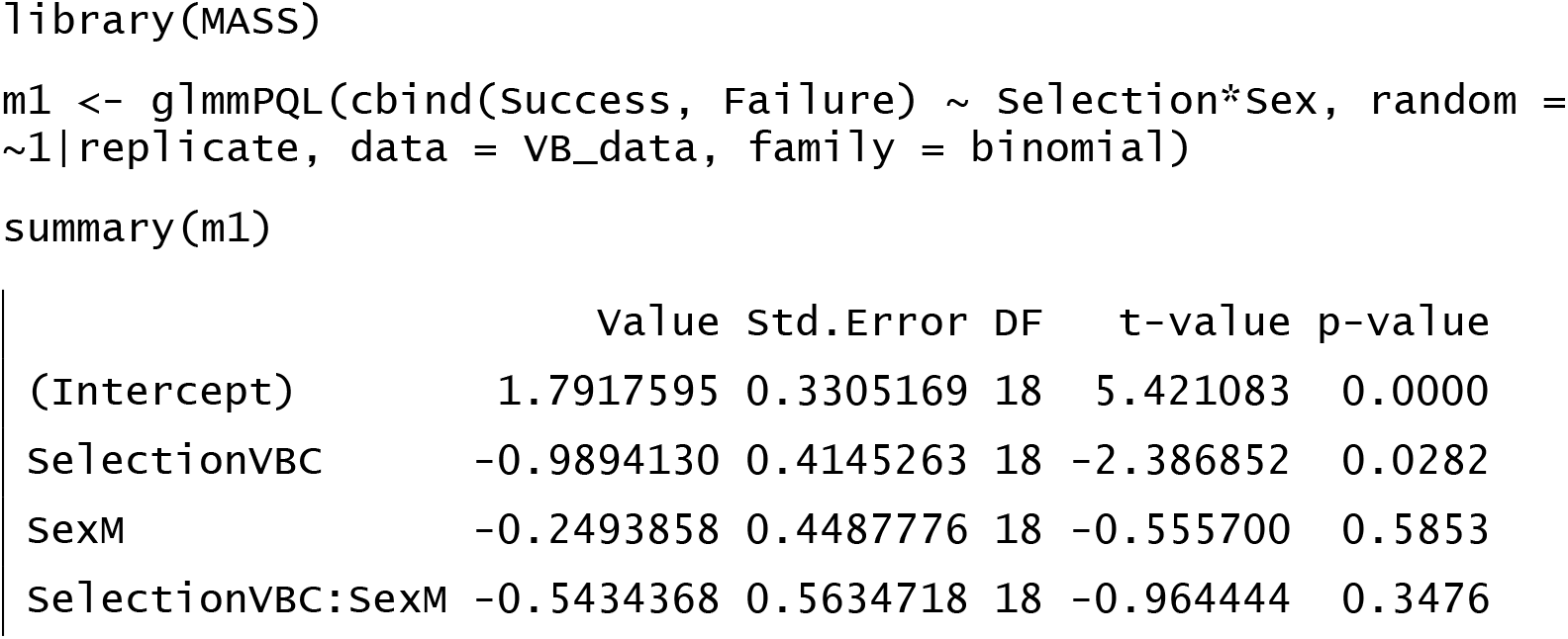

#### 2) Model reduction (removing the non-significant *dispersal selection × sex* interaction)

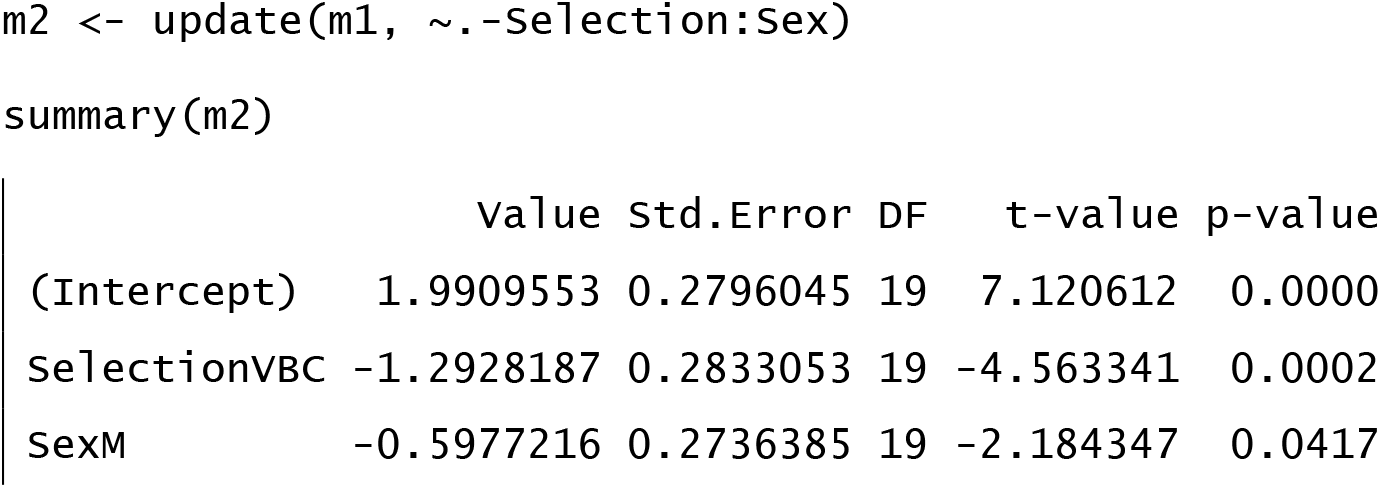

#### 3) Minimum Adequate Model (MAM): m2

### Text S2.4: Experiment 2 (DBS comparison)

#### 1) Full model

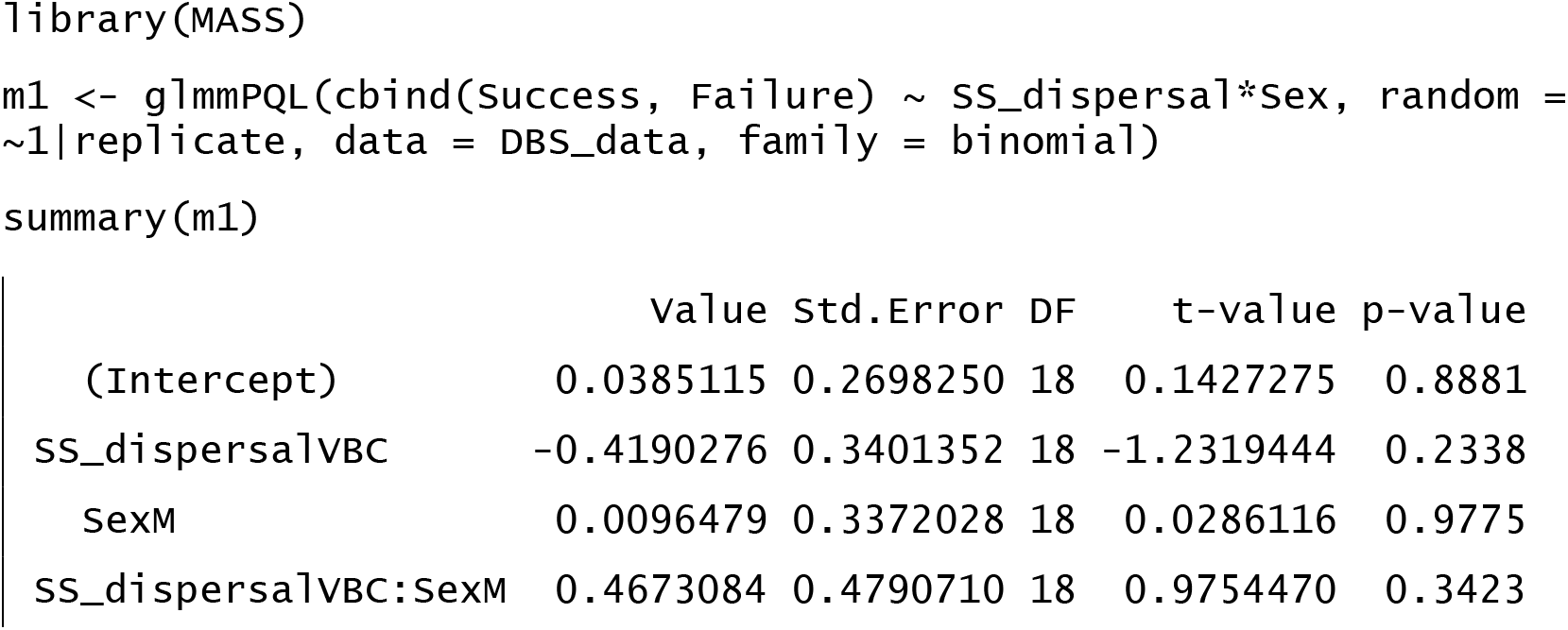

#### 2) Model reduction (removing the non-significant *same-sex dispersal × sex* interaction)

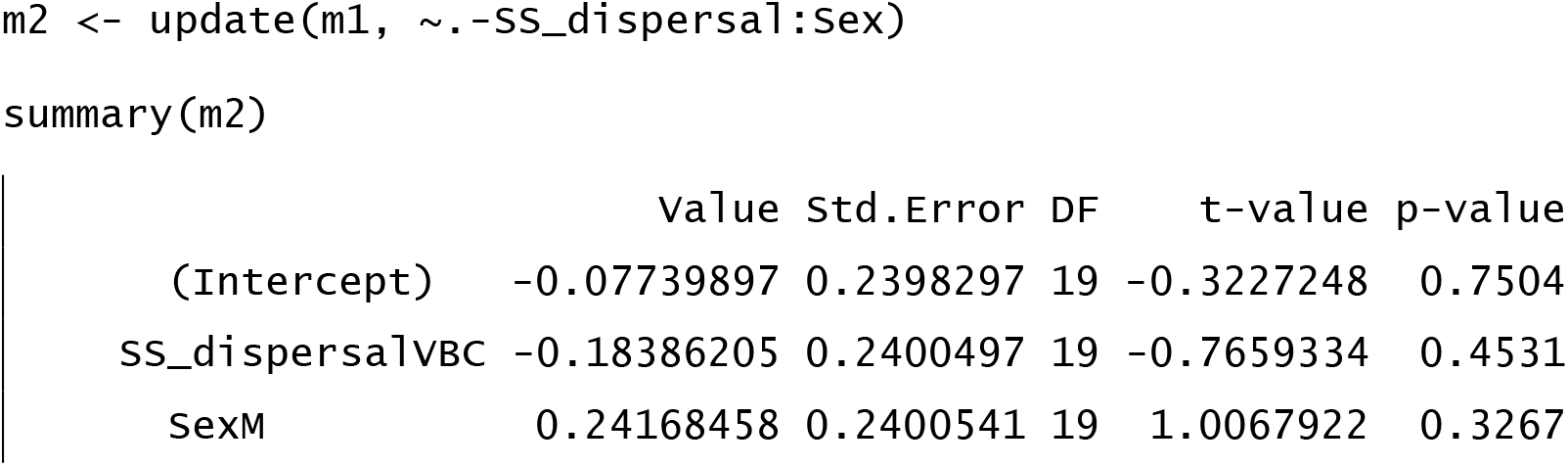

#### 3) Model reduction (removing the non-significant factor,*same-sex dispersal*)

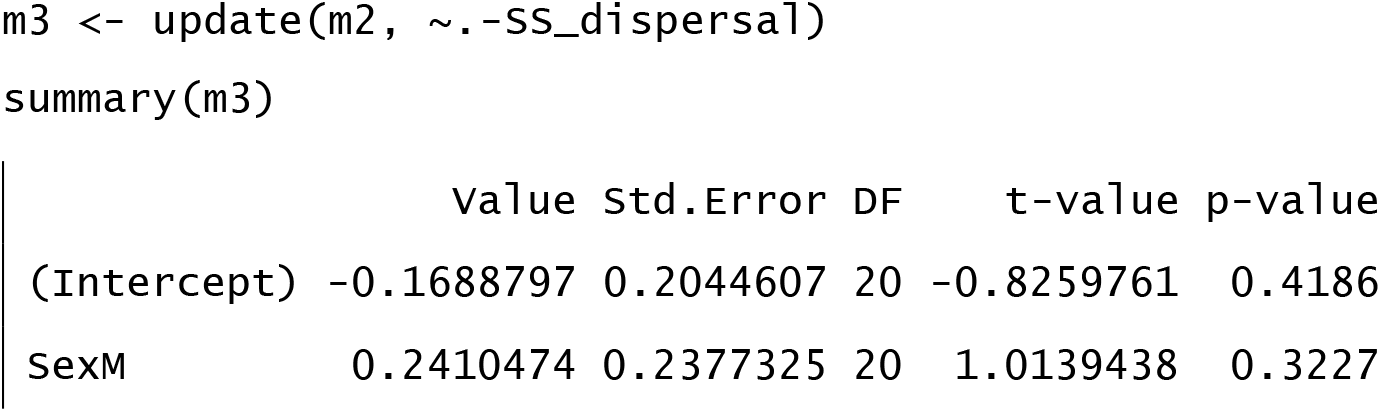

#### Minimum Adequate Model (MAM): m3

**Fig. S2.**
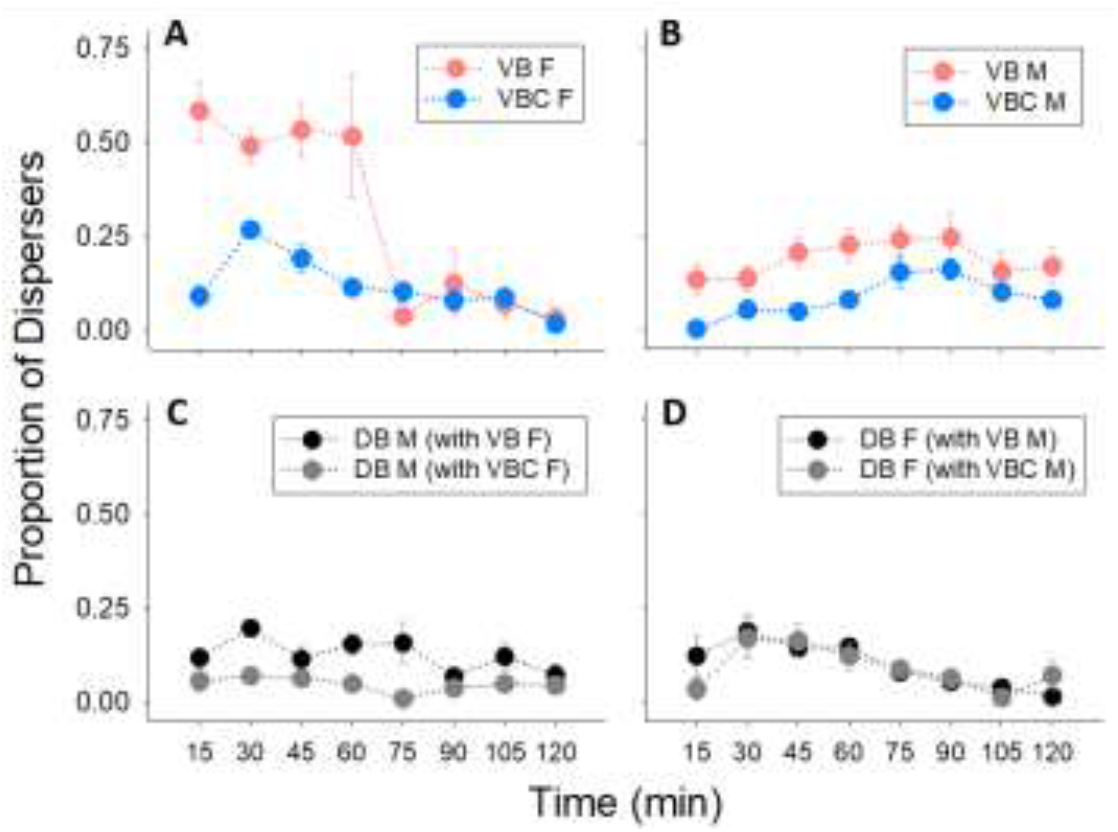
Temporal dispersal probability of individuals in the mixed-sex scenario (Experiment 1). Dispersers in each temporal bin plotted as the proportion of remaining (non-dispersed) individuals until that time point, for (A) VB (high-dispersive) vs. VBC (low-dispersive) females, (B) VB (high-dispersive) vs. VBC (low-dispersive) males, (C) DB (baseline) males dispersing with VB vs. VBC females, and (D) DB (baseline) females dispersing with VB vs. VBC males. Note that this is a different representation of the temporal dispersal profile presented in main Fig 3, where the proportion of dispersers sum to 1. Here, the proportion of remaining dispersers do not sum to any constant value.

**Fig. S3.**
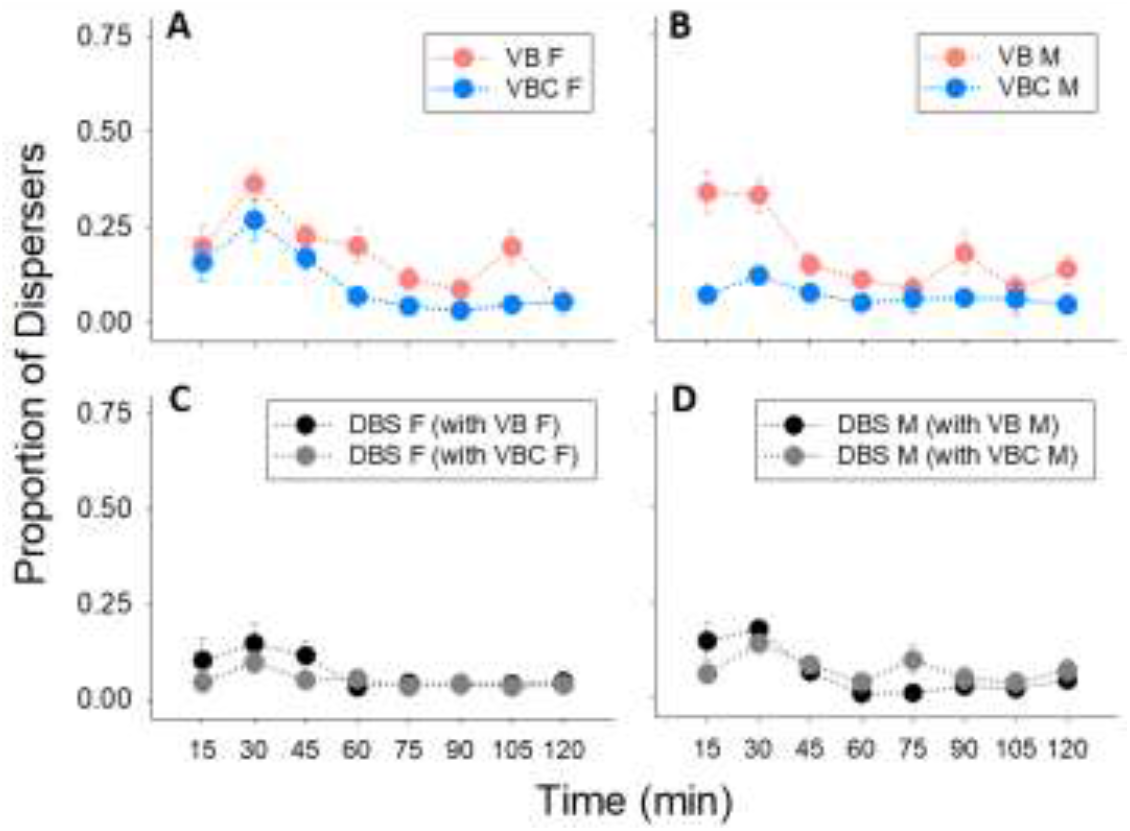
Temporal dispersal probability of individuals in the same-sex scenario (Experiment 2). Dispersers in each temporal bin plotted as the proportion of remaining (non-dispersed) individuals until that time point, for (A) VB (high-dispersive) vs. VBC (low-dispersive) females, (B) VB (high-dispersive) vs. VBC (low-dispersive) males, (C) DBS (scarlet-eyed baseline) females dispersing with VB vs. VBC females, and (D) DB (scarlet-eyed baseline) males dispersing with VB vs. VBC males. Note that this is a different representation of the temporal dispersal profile presented in main Fig 3, where the proportion of dispersers sum to 1. Here, the proportion of remaining dispersers do not sum to any constant value.

## Text S3: Model reduction details for temporal dispersal profile analyses

### Text S3.1: Experiment 1 (VB/VBC comparison)

#### 1) Full model

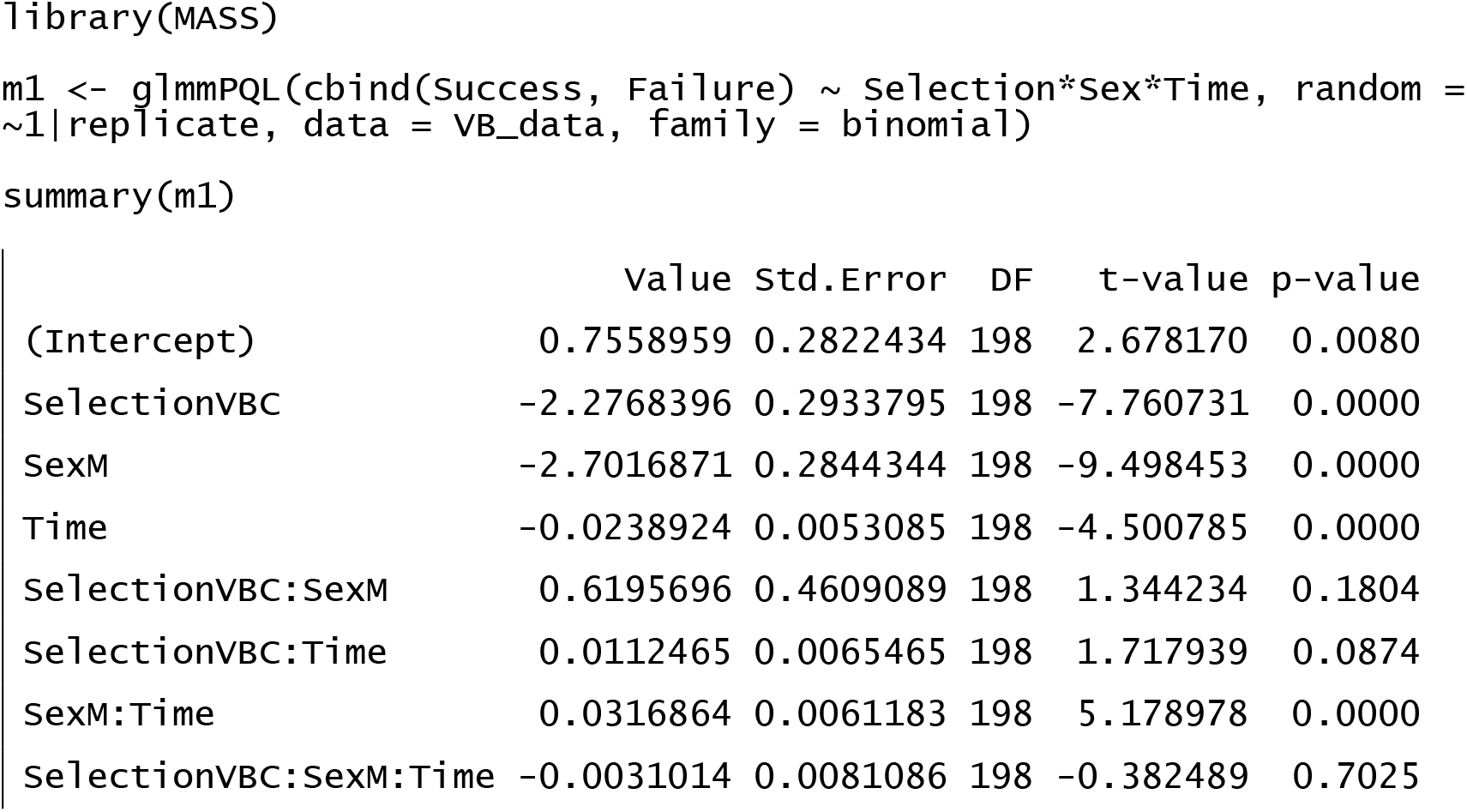

#### 2) Model reduction (removing the non-significant *dispersal selection × sex × time* interaction)

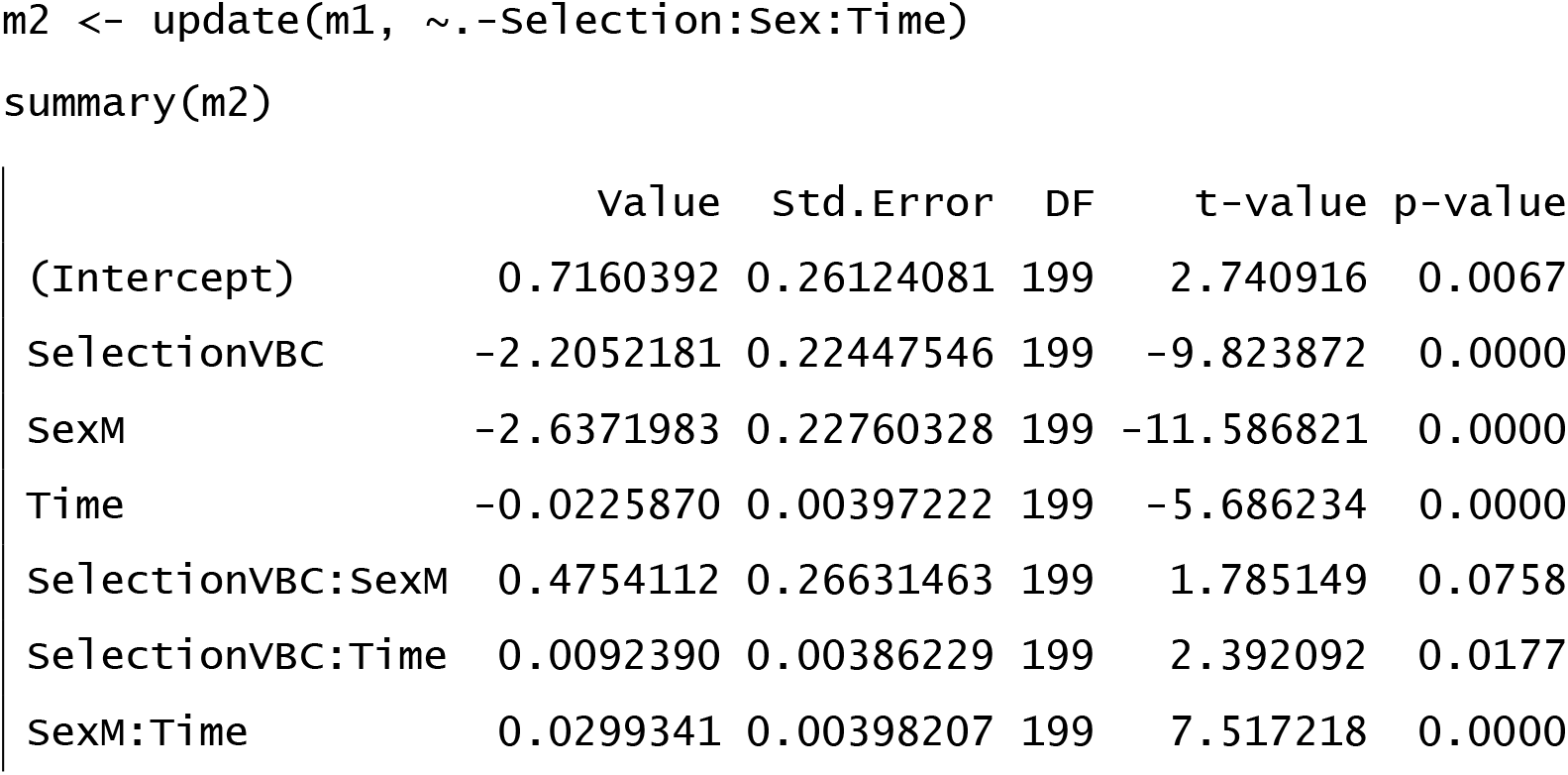

#### 3) Model reduction (removing the non-significant *dispersal selection × sex* interaction)

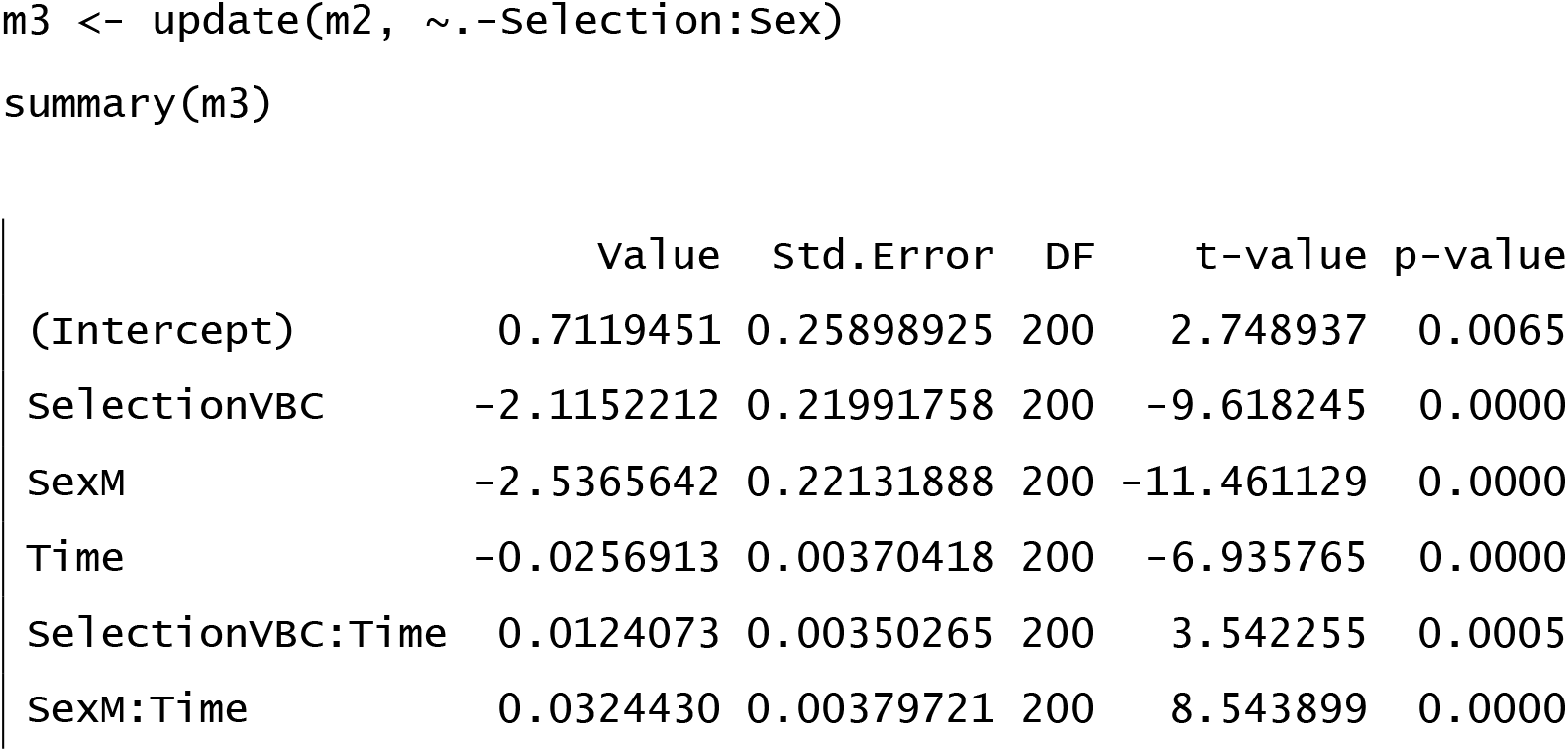

#### 4) Minimum Adequate Model (MAM): m3

### Text S3.2: Experiment 1 (DB comparison)

#### 1) Full model

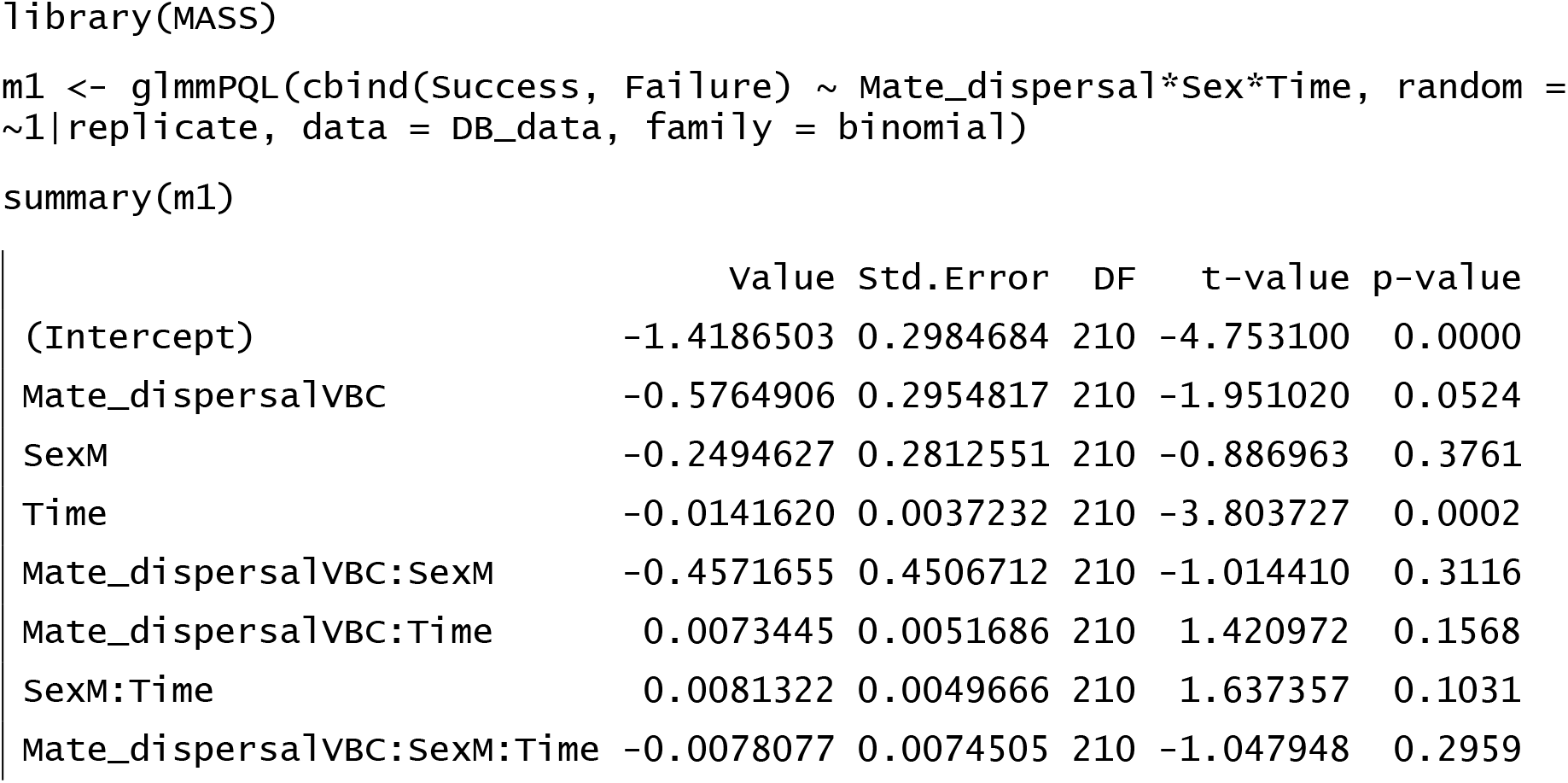

#### 2) Model reduction (removing the non-significant *mate dispersal × sex × time* interaction)

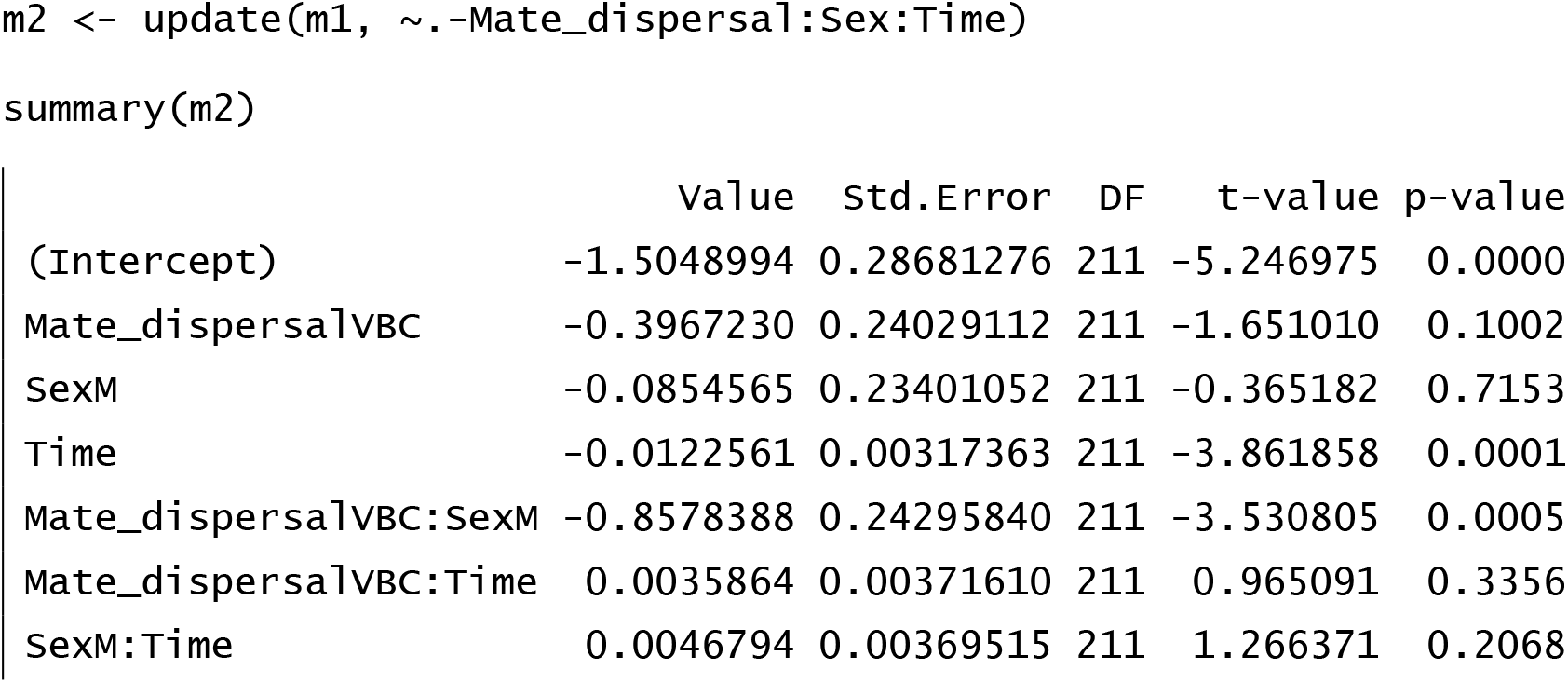

#### 3) Model reduction (removing the non-significant *mate dispersal × time* interaction)

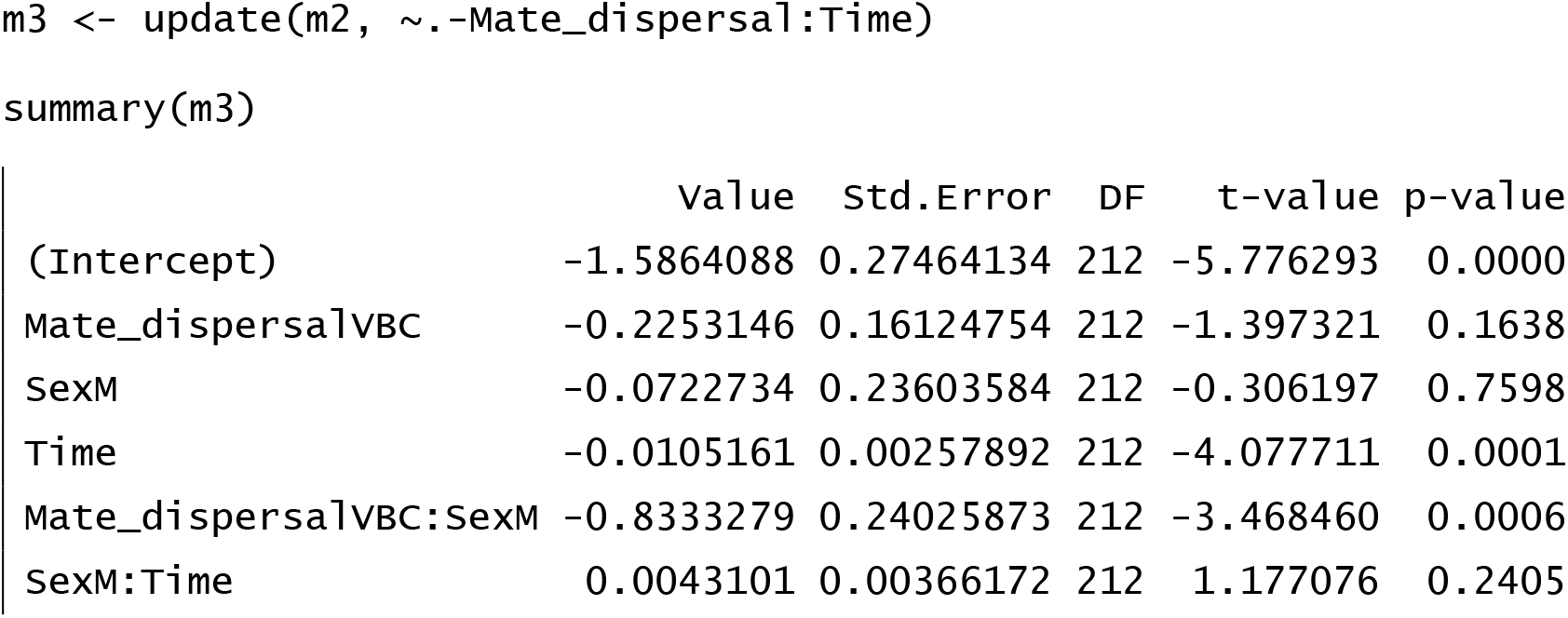

#### 4) Model reduction (removing the non-significant *sex × time* interaction)

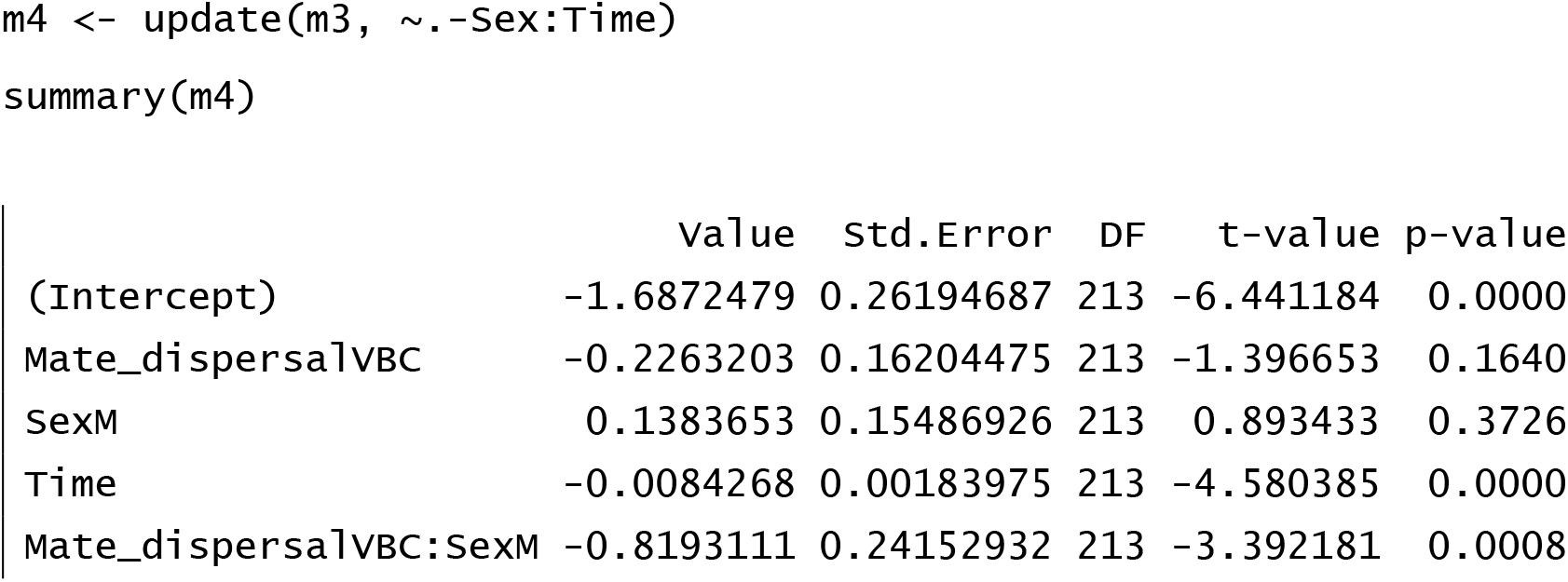

#### 5) Minimum Adequate Model (MAM): m4

### Text S3.3: Experiment 2 (VB/VBC comparison)

#### 1) Full model

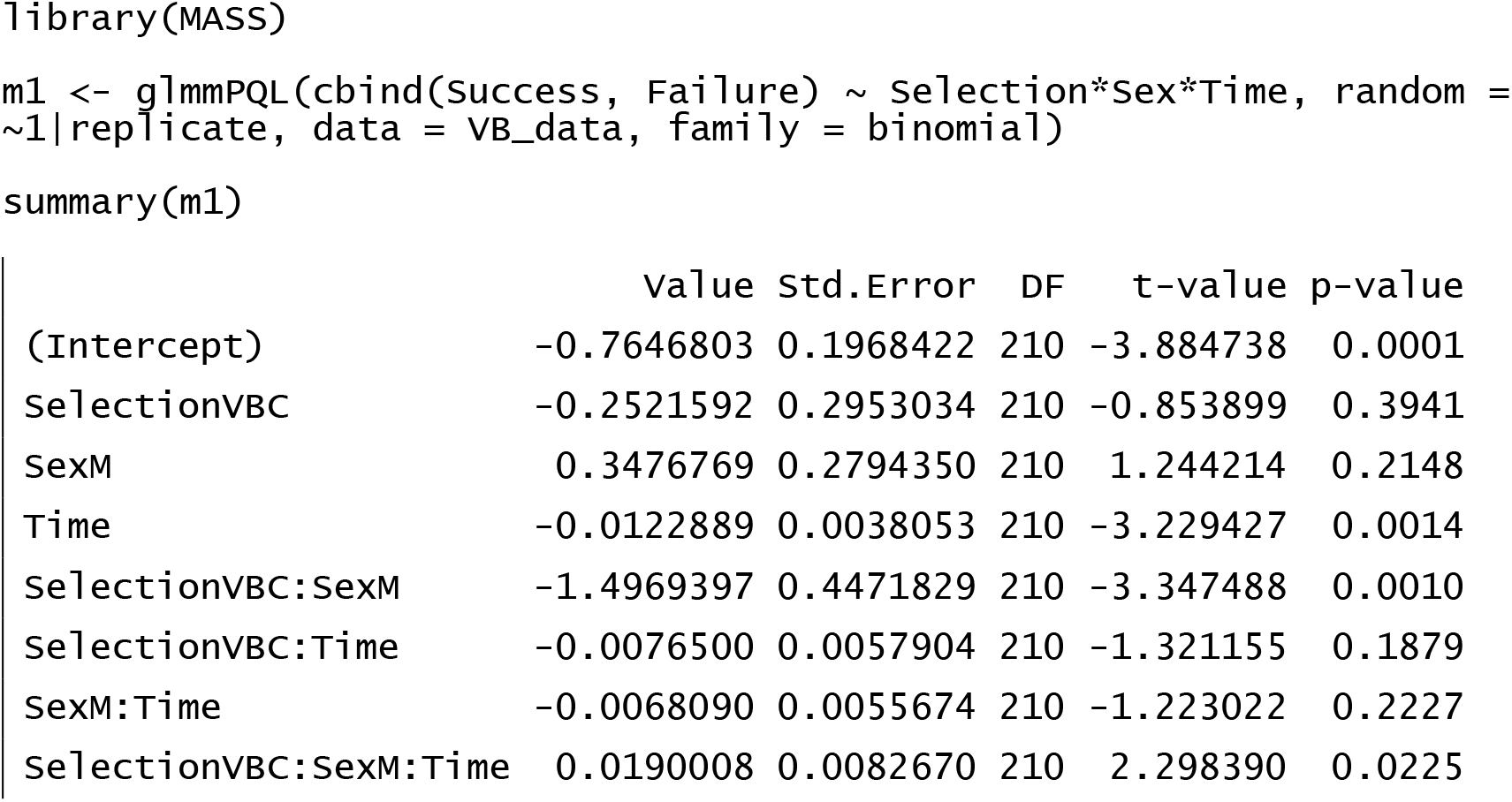

#### 2) Minimum Adequate Model (MAM): m1

### Text S3.4: Experiment 2 (DBS comparison)

#### 1) Full model

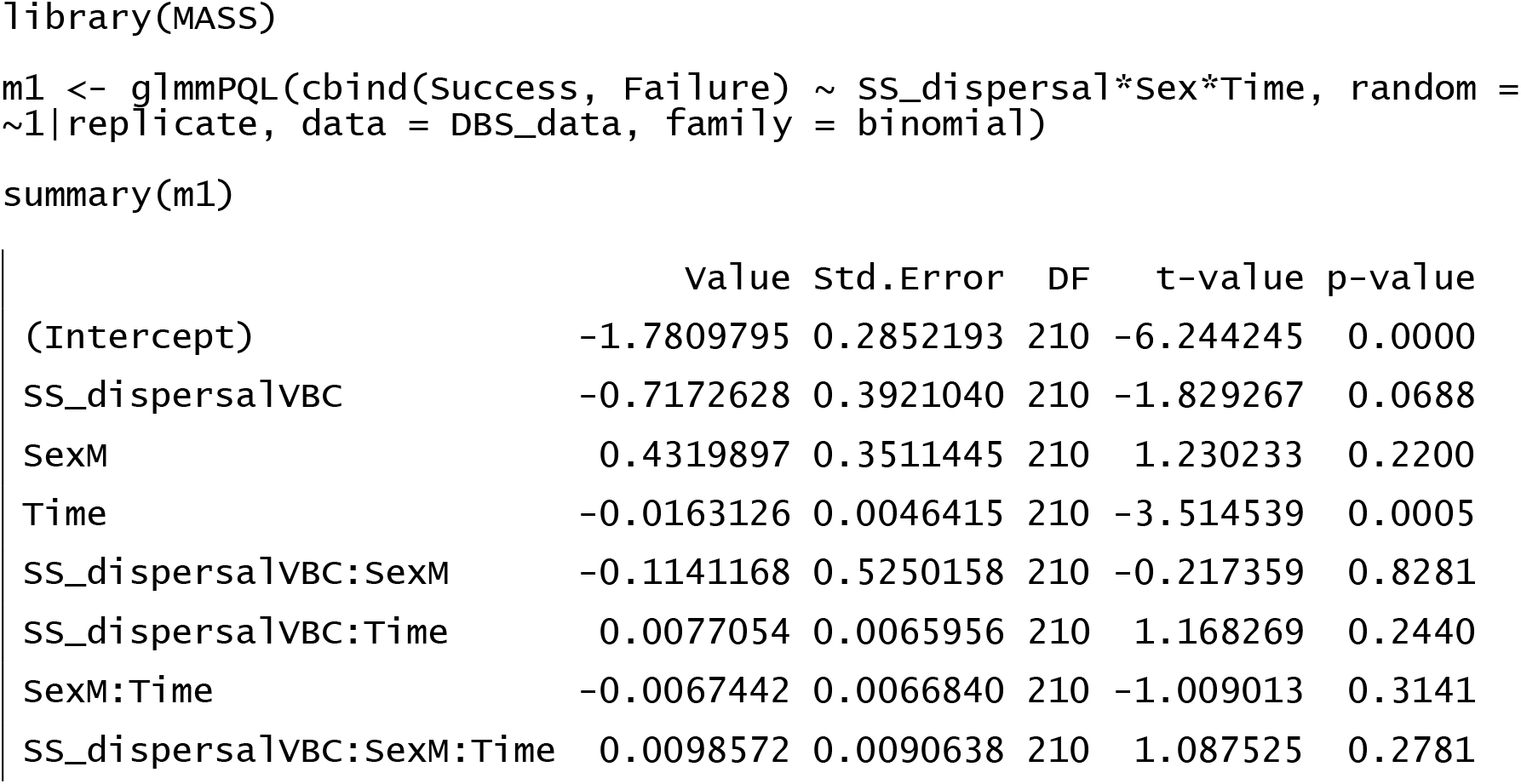

#### 2) Model reduction (removing the non-significant *same-sex dispersal × sex* × time interaction)

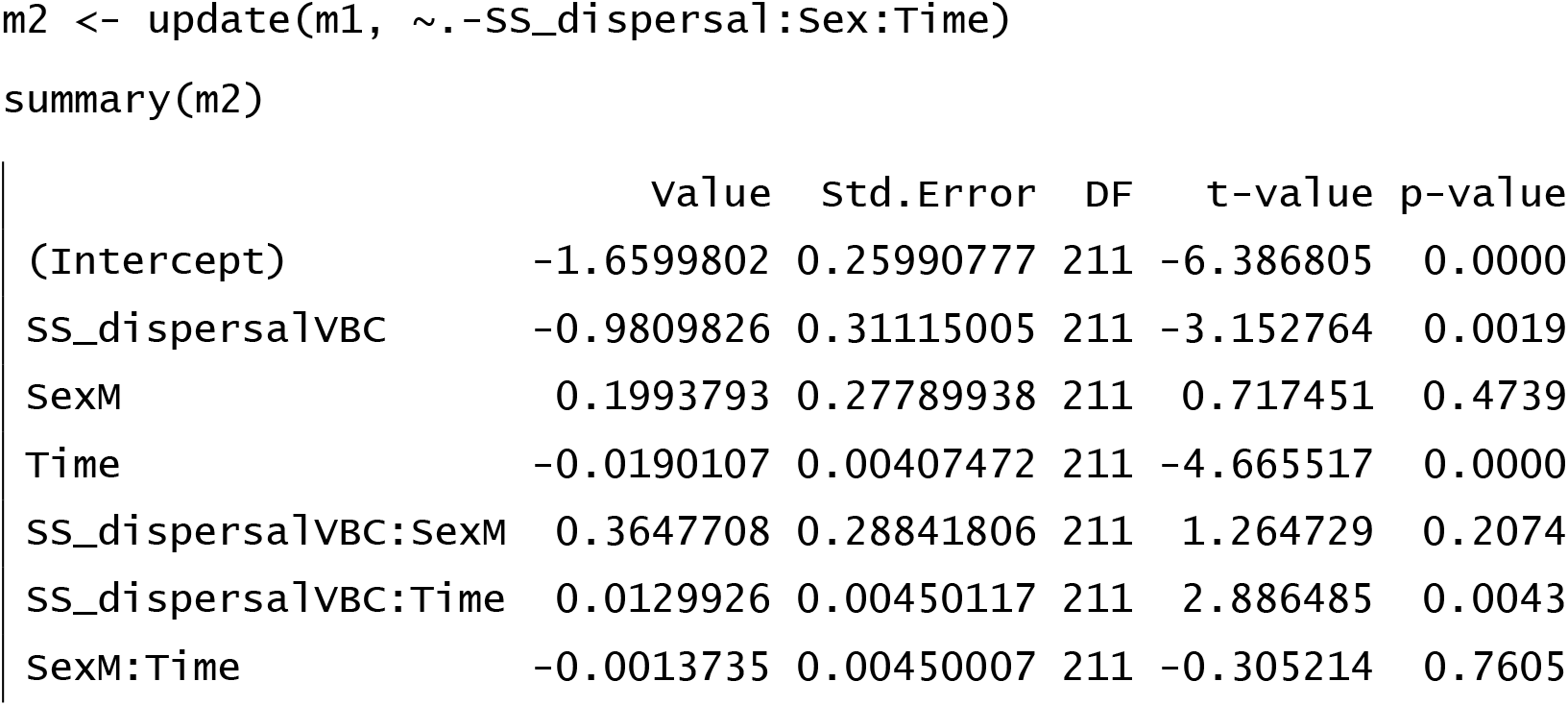

#### 3) Model reduction (removing the non-significant *sex × time* interaction)

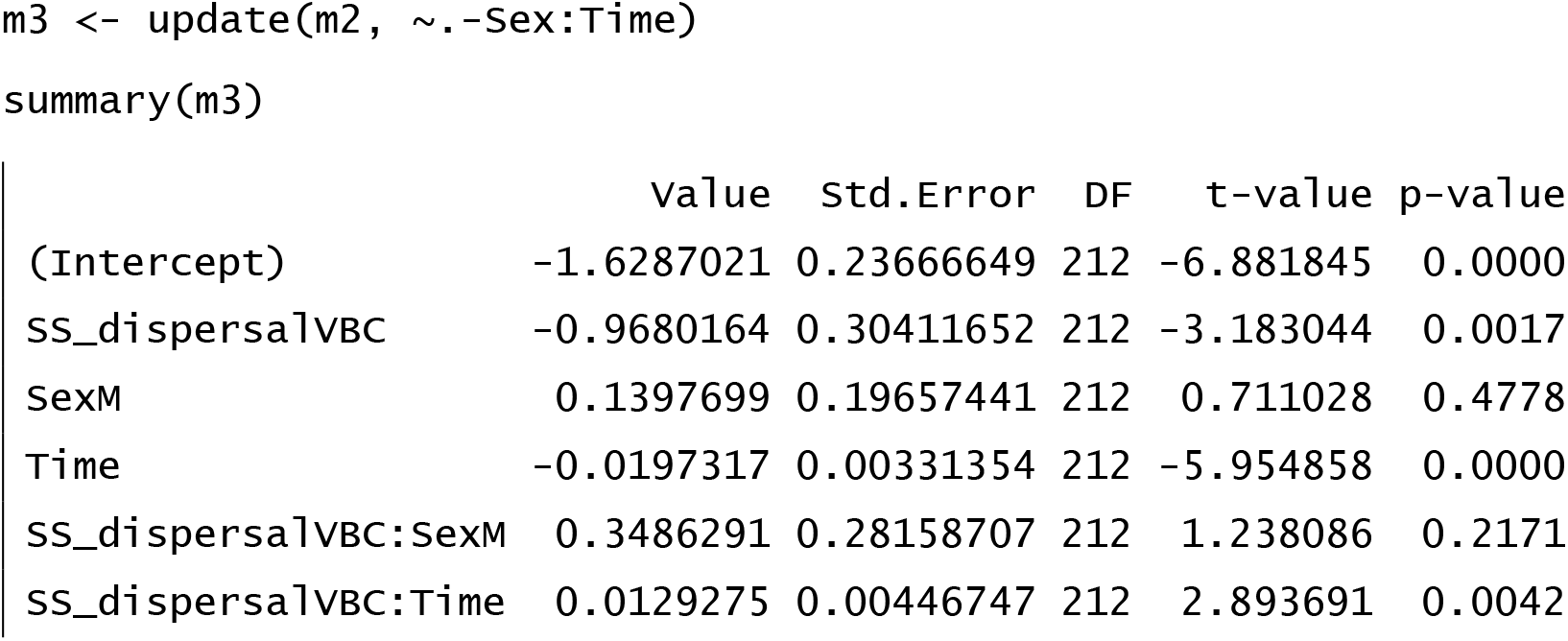

#### 4) Model reduction (removing the non-significant *same-sex dispersal × sex* interaction)

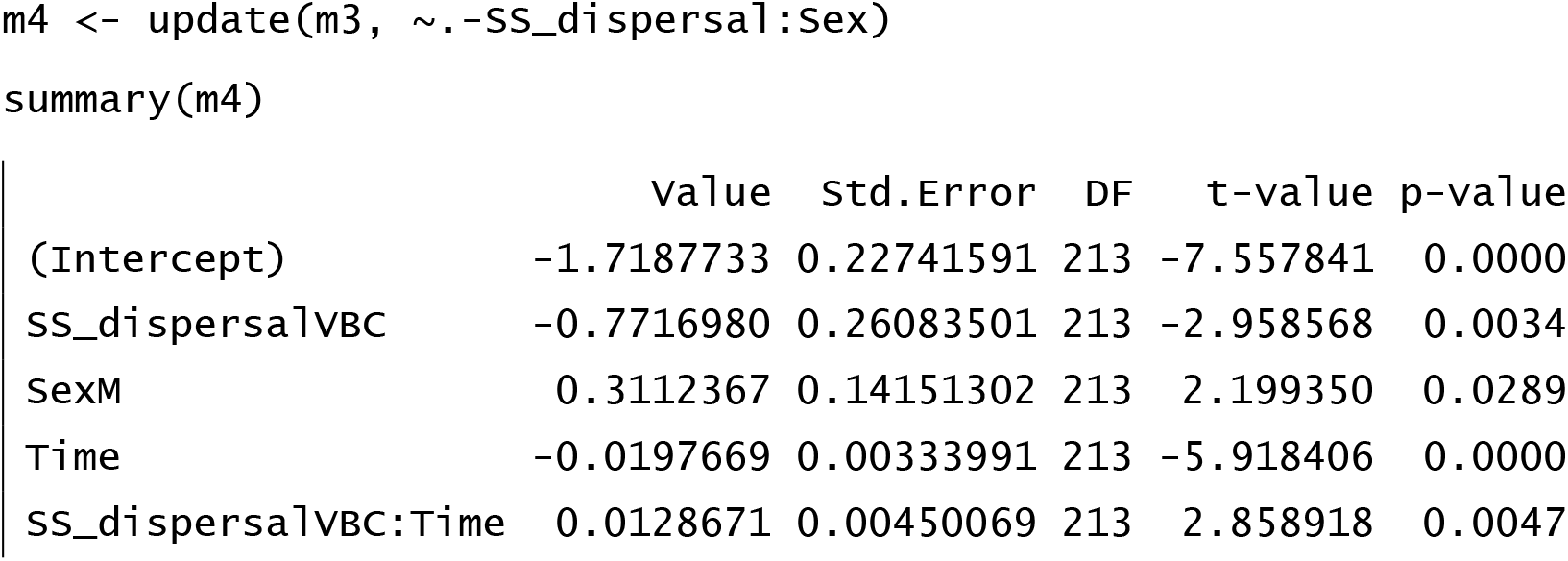

#### 5) Minimum Adequate Model (MAM): m4

